# The *Pseudomonas aeruginosa* lectin LecB causes integrin internalization to facilitate crawling of bacteria underneath host cells

**DOI:** 10.1101/2019.12.12.872739

**Authors:** Roland Thuenauer, Alessia Landi, Anne Trefzer, Silke Altmann, Sarah Wehrum, Thorsten Eierhoff, Britta Diedrich, Jörn Dengjel, Alexander Nyström, Anne Imberty, Winfried Römer

**Affiliations:** Faculty of Biology, Albert-Ludwigs-University Freiburg, Freiburg, Germany; BIOSS Centre for Biological Signalling Studies, Albert-Ludwigs-University Freiburg, Freiburg, Germany; Advanced Light and Fluorescence Microscopy Facility, Centre for Structural Systems Biology (CSSB), Hamburg, Germany; Department of Biology, University of Hamburg, Hamburg, Germany; Department of Biology, University of Fribourg, Fribourg, Switzerland; and Department of Dermatology, Medical Center - Albert-Ludwigs-University Freiburg, Freiburg, Germany; Department of Dermatology, Medical Center - Albert-Ludwigs-University Freiburg, Freiburg, Germany; Univ. Grenoble Alpes, CNRS, CERMAV, Grenoble, France

## Abstract

The opportunistic bacterium *Pseudomonas aeruginosa* produces the fucose-specific lectin LecB, which has been identified as virulence factor. LecB has a tetrameric structure with four opposing binding sites and has been shown to act as crosslinker. Here, we demonstrate that LecB strongly binds to the glycosylated moieties of β1-integrins on the basolateral plasma membrane of epithelial cells and caused rapid integrin endocytosis. Whereas internalized integrins were degraded via a lysosomal pathway, washout of LecB restored integrin cell surface localization, thus indicating a specific and direct action of LecB on integrins to bring about their endocytosis. Interestingly, LecB was able to trigger uptake of active and inactive β1-integrins and also of complete α3β1-integrin - laminin complexes. We provide a mechanistic explanation for this unique endocytic process by showing that LecB has the additional ability to recognize fucose-bearing glycosphingolipids and caused the formation of membrane invaginations on giant unilamellar vesicles. In cells, LecB recruited integrins to these invaginations by crosslinking integrins and glycosphingolipids. In epithelial wound healing assays, LecB specifically cleared integrins from the surface of cells located at the wound edge and blocked cell migration and wound healing in a dose-dependent manner. Moreover, the wild type *P. aeruginosa* strain PAO1 was able to loosen cell-substrate adhesion in order to crawl underneath exposed cells, whereas knockout of LecB significantly reduced crawling events. Based on these results we suggest that LecB has a role in disseminating bacteria along the cell - basement membrane interface.

**Importance:** *Pseudomonas aeruginosa* is a ubiquitous environmental bacterium that is one of the leading causes for nosocomial infections. *P. aeruginosa* is able to switch between planktonic, intracellular, and biofilm-based lifestyles, which allows it to evade the immune system as well as antibiotic treatment. Hence, alternatives to antibiotic treatment are urgently required to combat *P. aeruginosa* infections. Lectins, like the fucose-specific LecB, are promising targets, because removal of LecB resulted in decreased virulence in mouse models. Currently, several research groups are developing LecB inhibitors. However, the role of LecB in host-pathogen interaction is not well understood. The significance of our research is in identifying cellular mechanisms how LecB facilitates *P. aeruginosa* infection: We introduce LecB as new member to the list of bacterial molecules that bind integrins and show that *P. aeruginosa* can efficiently move forward underneath attached epithelial cells by loosening cell - basement membrane attachment in a LecB-dependent manner.

## 1 Introduction

*Pseudomonas aeruginosa* is a ubiquitous Gram-negative environmental bacterium. For humans it acts as an opportunistic pathogen and can cause severe infections, predominantly in cystic fibrosis patients (1) and immunocompromised individuals, such as HIV patients (2), patients receiving cancer treatment (3), patients with assisted ventilation (4), and patients with burn wounds (5). *P. aeruginosa* infections are difficult-to-treat because the bacterium has a high natural resistance to antibiotics and rapidly acquires new antibiotic resistances (6). In fact, several outbreaks caused by multidrug-resistant *P. aeruginosa* strains were recently reported (7, 8). In addition, the bacterium is able to adopt various lifestyles that allow it to evade the immune system as well as antibiotic treatment. In particular, *P. aeruginosa* can form biofilms (9) and invades and proliferates in host cells (10). These properties make *P. aeruginosa* an imminent threat for global health and therefore the World Health Organization (WHO) categorized *P. aeruginosa* with ‘Priority 1’ on its recently released ‘WHO priority pathogens list for research and development of new antibiotics’ (11), which highlights the need to develop novel treatment strategies for *P. aeruginosa* infections (12).

When infecting the human body, *P. aeruginosa* typically encounters polarized epithelial cell layers, which function as protective barriers (10). As an opportunistic bacterium, *P. aeruginosa* adapts its strategy according to the circumstances it encounters. It harnesses weak spots, for example sites where cells divide or are extruded, to proceed to the basolateral side of epithelia (13). *P. aeruginosa* has also been shown to have a propensity to enter and colonize wounded epithelia (10), and there is ample experimental evidence that loss of epithelial polarity increases detrimental effects of *P. aeruginosa* on host cells (10). In addition, *P. aeruginosa* has evolved strategies to manipulate the polarity of host epithelial cells to facilitate infection (10, 14). When reaching the basolateral side, *P. aeruginosa* gets access to integrins, which are typically restricted to the basolateral plasma membrane of epithelial cells. Although integrins are well known as receptors for multiple pathogens (15–17), and previous studies have shown that *P. aeruginosa* is able to bind to α5β1-integrins in nasal epithelial cells (18) and to αvβ5-integrins in lung epithelial cells (19), the specific roles for integrins for *P. aeruginosa* infection remain unclear.

*P. aeruginosa* produces two carbohydrate-binding proteins, so-called lectins, LecA and LecB, which are also named PA-IL and PA-IIL, respectively (20). Whereas LecA is galactophilic, LecB prefers fucose (20). LecB is transported to the outer bacterial membrane, where it binds to the porin OprF, resulting in its presentation at the outer surface of *P. aeruginosa* (21, 22). Several lines of evidence indicate that LecB is an important virulence factor. LecB-deficient *P. aeruginosa* are less pathogenic (23) and show diminished biofilm formation (21). In addition, LecB was found to abrogate ciliary beating in human airways (24) and to diminish tissue repair processes in lung epithelia (25). These findings raised the prospect to establish alternative treatment strategies for *P. aeruginosa* infections by blocking LecB, and stimulated ongoing efforts of several research groups to develop LecB inhibitors (26–31).

However, the functions of LecB remain difficult to pin down, because as a lectin it can bind to many different host cell receptors. Here, we demonstrate that integrins are major receptors of LecB. Moreover, we observed that LecB binding to integrins resulted in their rapid cellular uptake together with their basement membrane ligands. We provide a mechanistic explanation for this distinctive endocytosis process by showing that LecB binding to fucose-bearing lipids induces membrane invaginations and, furthermore, LecB positions integrins in these invaginations by crosslinking integrins and lipids. As functional consequence, purified LecB caused inhibition of cell migration and abrogation of epithelial wound healing by specifically internalizing exposed integrins in cells at the wound edge. Furthermore, we could demonstrate that the wild type (wt) *P. aeruginosa* strain PAO1 is able to locally disturb cell adhesion and to crawl underneath epithelial cells. Importantly, knocking out LecB diminished the number of *P. aeruginosa* found underneath epithelial cells, thus implicating LecB as virulence factor enabling bacteria to colonize host tissue along the interface between cells and the basement membrane.

## 2 Results

### 2.1 Differential effects of LecB at the apical and basolateral side of polarized epithelial cells

When *P. aeruginosa* infects a human body, it typically encounters first the apical pole of epithelial cells. Through induced or pre-existing damages, the bacterium can access the basolateral cell pole of epithelial cells. Since the apical and basolateral plasma membrane of individual epithelial cells harbors distinct sets of membrane proteins and lipids, we investigated if LecB causes different effects when applied to the apical or basolateral side. We chose Madin-Darby canine kidney (MDCK) cells as model system because they reliably form polarized monolayers *in vitro* (32, 33) and have been already successfully used in *P. aeruginosa* infection studies (14, 34). Purified LecB was able to bind apical and basolateral plasma membranes of MDCK cells (Fig. S1A). Interestingly, apical application of LecB resulted in completely different responses of the host cells than basolateral application (Fig. 1A). After 6 h and 12 h of apical treatment with 50 μg/ml (4.3 μM) LecB the overall morphology of the cells was intact as evidenced by staining of β-catenin (red) that remained basolateral and GPI-GFP (green) that remained apical. In addition, tight junction integrity was not disturbed as demonstrated by the unchanged staining of ZO-1 (white in Fig. 1A) and the preserved trans-epithelial electrical resistance (TEER; Fig. 1B). In contrast, 6 h and 12 h of basolateral treatment with 50 μg/ml LecB resulted in rounded cell morphologies and severely disturbed epithelial polarity. GPI-GFP became localized all around the cells and tight junctions almost disappeared (Fig. 1A), which was corroborated by a drastic reduction of the TEER (Fig. 1B). Importantly, these effects cannot be explained by potential LecB-mediated apoptosis or necrosis (Fig. S1B-C). Yet, the observed consequences seem to be specific for LecB, because another fucose-binding lectin, *Ulex europaeus* agglutinin I (UEA-I (35)), which did also bind to apical and basolateral plasma membranes of MDCK cells (Fig. S1D), did not cause apparent changes in cell morphology (Fig. S1E), nor did it influence the TEER (Fig. S1F).

**Fig. 1:**
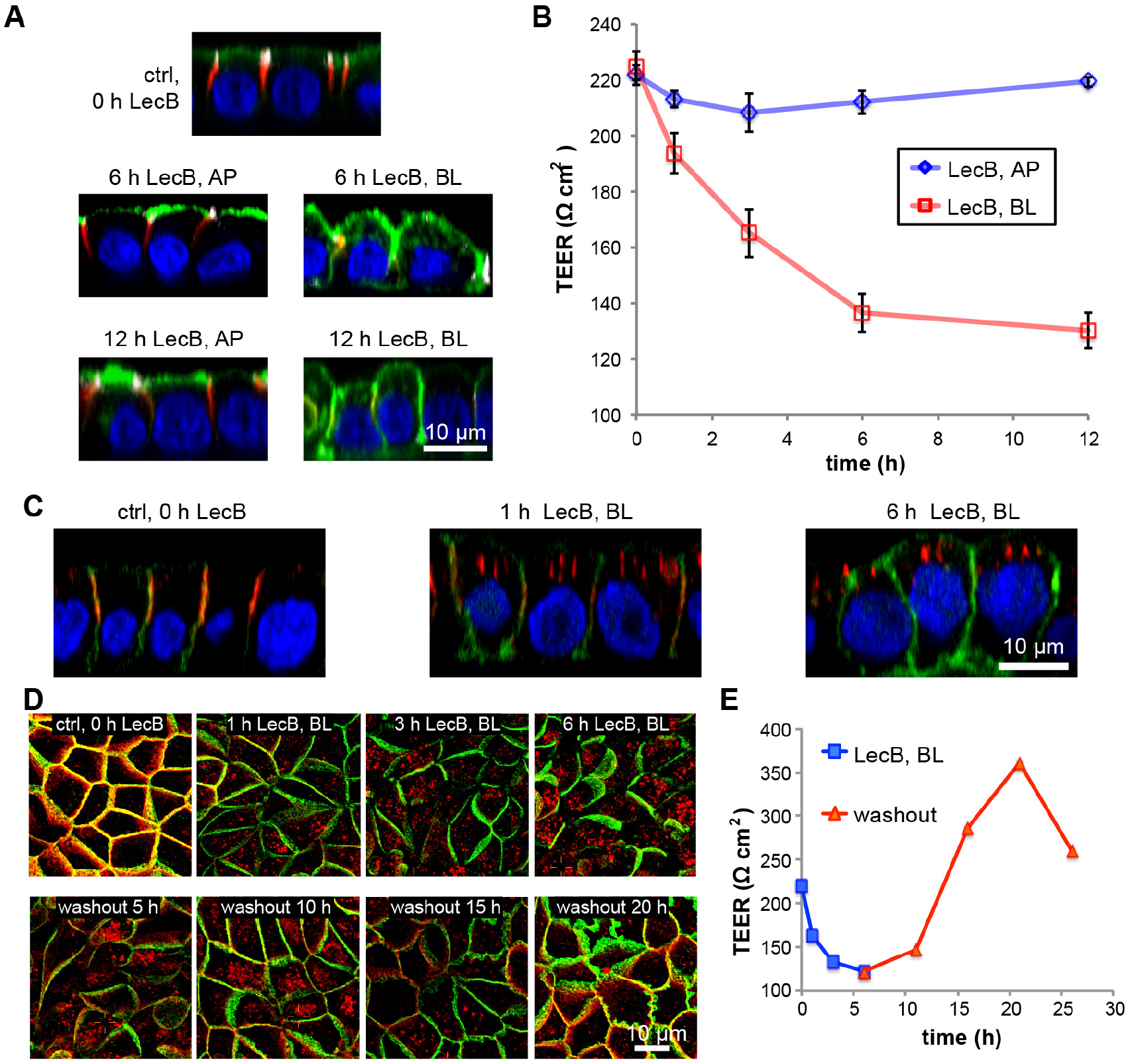
Basolateral LecB application depolarized MDCK cells and caused integrin internalization. (A) Polarized, filter-grown MDCK cells stably expressing the apical marker GPI-GFP (green) were left untreated (ctrl) or treated apically (AP) or basolaterally (BL) with 50 μg/ml LecB for the indicated time periods, fixed and stained with antibodies recognizing the basolateral marker β-catenin (red) and the tight junction marker ZO-1 (white), nuclei were stained with DAPI (blue). Representative sections along the apico-basal axis (x-z sections) extracted from confocal image stacks are shown. (B) Time course of the trans-epithelial electrical resistance (TEER) of MDCK monolayers treated AP or BL with LecB. The mean values from n = 3 experiments are displayed. (C) LecB was applied BL to MDCK cells stably expressing PH-Akt-GFP (green) for the indicated time periods. Cells were fixed and stained for β1-integrin (red); nuclei were stained with DAPI (blue). Representative x-z sections extracted from confocal image stacks are depicted. (D) MDCK cells were treated with LecB as indicated, fixed, and stained for β1-integrin (red) and β-catenin (green). Maximum intensity projections of confocal image stacks covering total cell heights are shown. (E) The time course of the TEER of MDCK cells treated BL with LecB as indicated and after washout was measured.

Taken together, basolateral application of LecB dissolves epithelial polarity in MDCK cells, whereas another fucose-binding lectin, UEA-I, does not cause such effects.

### 2.2 Basolaterally applied LecB binds β1-integrin and causes its internalization

To uncover the mechanisms of LecB-induced loss of epithelial polarity, we monitored the localization of cell adhesion receptors upon basolateral LecB stimulation. This revealed a rapid and efficient internalization of β1-integrins (Fig. 1C). Interestingly, this effect was reversible after washout of LecB after 6 h (Fig. 1D), and the timing of β1-integrin internalization and return to the cell surface correlated well with decrease and increase of the TEER (Fig. 1E).

To elucidate LecB-triggered β1-integrin internalization, we first investigated if LecB binds to β1-integrin. To this end we used LecB-biotin to precipitate LecB-receptor complexes with streptavidin beads. Western blot analysis of the precipitates showed that LecB-biotin is able to bind to β1-integrins only when applied to the basolateral side (Fig. 2A). The binding of LecB-biotin to β1-integrin appeared to be strong, since LecB-biotin was able to extract approximately 75% of total β1-integrin when applied to the basolateral side, as quantified from the band intensities of the Western blot. In addition, fluorescently labeled LecB co-localized with internalized β1-integrins (Fig. 2B) and was able to bind immunoprecipitated β1-integrin in a far-Western assay in dependence of the β1-integrin glycosylation status (Fig. 2C), which provides complementary evidence for the capacity of LecB to bind to glycosylated β1-integrin.

**Figure 2:**
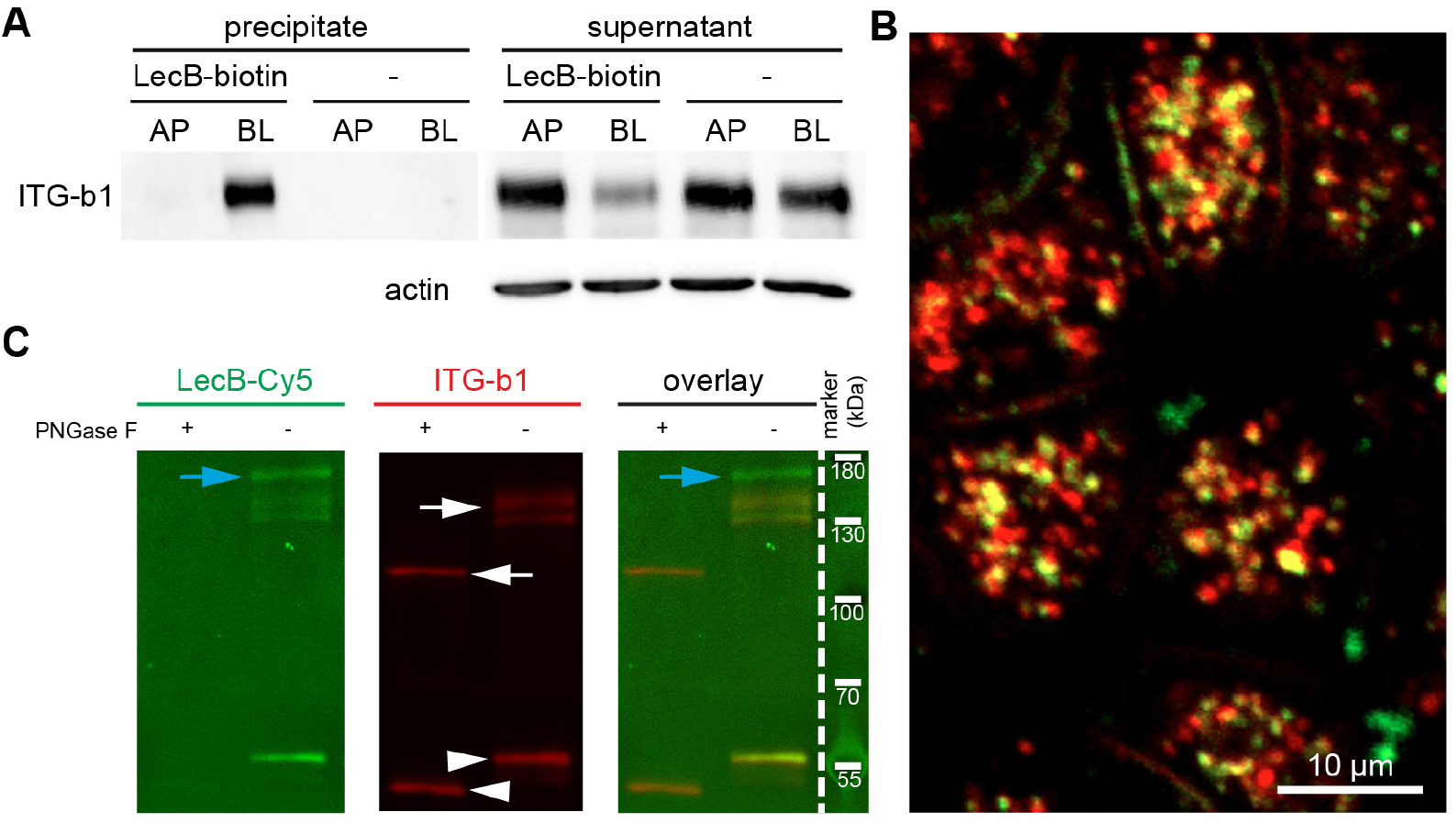
LecB directly binds to β1-integrin. (A) LecB-biotin was applied apically (AP) or basolaterally (BL) to polarized filter grown MDCK cells or cells were mock-treated AP or BL. Cells were lysed and LecB-biotin-receptor complexes were precipitated with streptavidin beads. Afterwards the presence of β1-integrin was probed by Western blot in the precipitate and the remaining supernatant of the precipitation. (B) LecB-Cy3 (red) was applied basolaterally to MDCK cells for 6 h. Cells were fixed and stained for β1-integrin (green). A confocal section (x-y section) crossing the cells in the sub-apical region is displayed, since in this region most internalized vesicles were concentrated. (C) MDCK cells were lysed, β1-integrins were immunoprecipitated and treated or left untreated with PNGase F to remove N-linked glycans. Western blot analysis of the immunoprecipitated β1-integrins was performed and β1-integrin presence was proven by staining with anti-β1-integrin antibodies (white arrows). Also bands from the antibody used for β1-integrin precipitation are visible (white arrowheads) and proteins that putatively co-precipitated with β1-integrin (blue arrows). To probe the binding of LecB to β1-integrin, LecB-Cy5 was incubated with membranes (far Western assay).

In summary, LecB recognizes β1-integrin at the basolateral cell surface and causes its rapid internalization.

### 2.3 α3-integrin and laminin are also internalized and degraded upon basolateral LecB application

From the far-Western assay in Fig. 2C it can be seen that LecB recognized not only glycosylated β1-integrin, but also other receptors that were presumably co-precipitated during β1-integrin immunoprecipitation (blue arrows). Hence, in a next step we identified basolateral interaction partners of LecB by LecB-biotin co-precipitation followed by mass spectrometry analysis and found 65 profoundly enriched proteins (Table S1). This analysis revealed that LecB is able to pull down virtually all integrins expressed by MDCK cells (36), and also many proteins known to interact with integrins, such as tetraspanins, basigin, EGFR, were detected. From this it appears that integrins are major cellular receptors of LecB. We focused our further analysis on α3β1-integrin and were able to demonstrate that α3-integrins are co-internalized with β1-integrins upon basolateral LecB application (Fig. 3A). Surprisingly, also the major ligands of α3β1-integrin expressed by MDCK cells, laminin-332 and/or −511 (37), were co-internalized (Fig. 3B). This suggests that LecB is able to cause endocytosis of intact α3β1-integrin – laminin complexes. To measure the dynamics of α3β1-integrin internalization we carried out surface biotinylation experiments (38). These experiments confirmed the rapid LecB-triggered internalization of α3- and β1-integrin subunits (Figs. 3C – 3E). In addition, the surface biotinylation experiments revealed that also the intracellular amount of α3- and β1-integrin subunits decreased upon LecB stimulation, suggesting a degradation of internalized integrins. Consistently, LecB-mediated reduction of integrins was also detected when whole cell lysates were subjected to Western blot analysis (Fig. S2A). Loss of integrins by degradation is further supported by our finding that internalized integrins after basolateral LecB treatment showed a time-dependent increase in co-localization with the late endosome marker Rab9 (Figs. S2B-C) and the lysosome marker Lamp1 (Figs. S2D-E).

**Figure 3:**
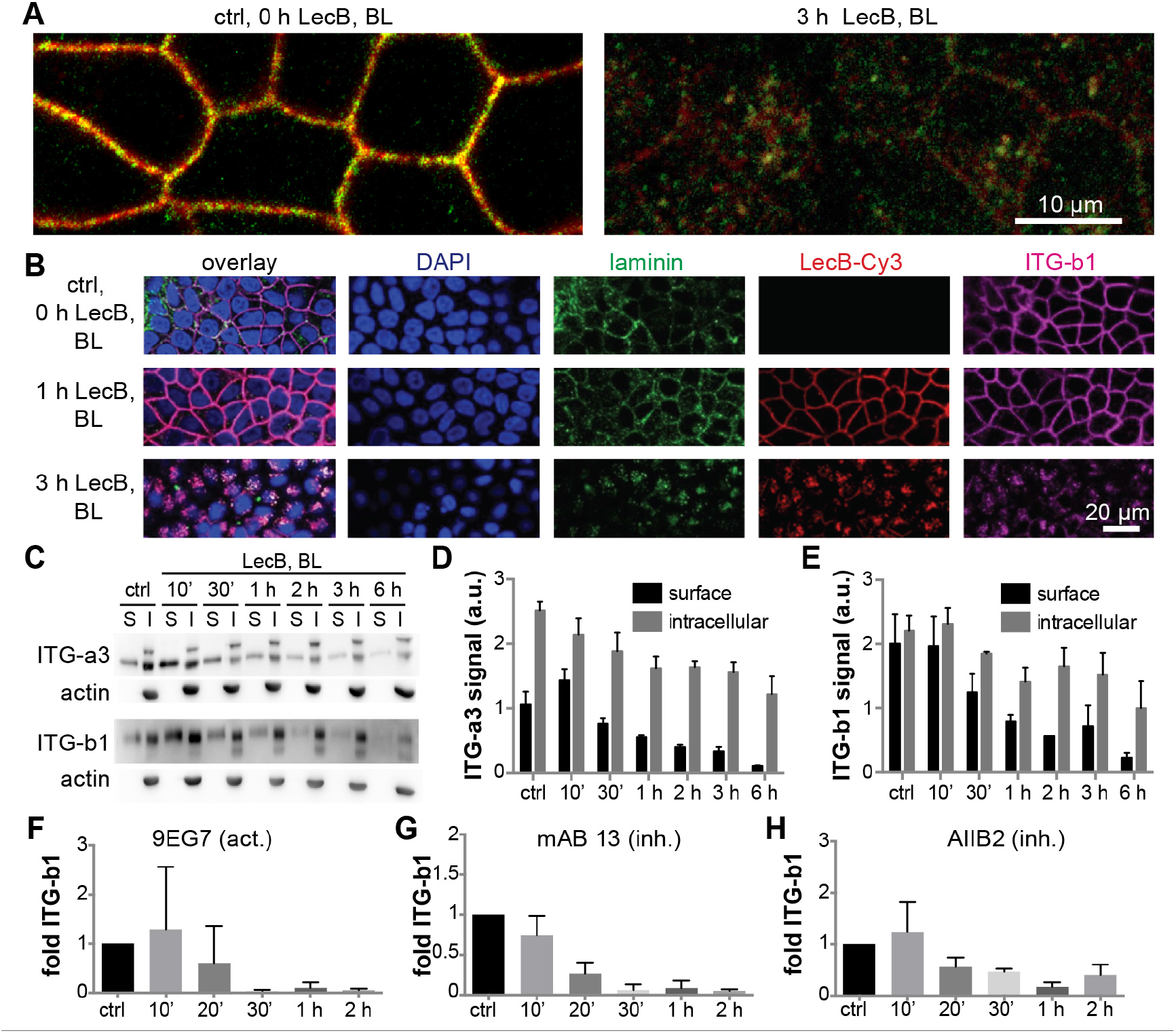
LecB internalizes α3β1-integrin regardless of its activation status and together with laminins. (A) MDCK cells were treated with LecB as indicated, fixed with methanol, and stained for α3-integrin (red) and β1-integrin (green). Confocal sections (x-y sections) through the middle of the cells extracted from confocal image stacks are shown. (B) MDCK cells were treated with LecB-Cy3 (red) as indicated, fixed, and stained for pan-laminin (green), and β1-integrin (magenta); nuclei were stained with DAPI (blue); x-y confocal sections through the middle of the cells are depicted. (C) – (E) MDCK cells were treated with LecB as indicated and surface biotinylation from the basolateral side was performed. After precipitation of biotinylated proteins, the precipitates representing the surface fraction (S) and the supernatant representing the intracellular fraction (I) were subjected to Western blot analysis and α3-integrins and β1-integrins were probed, as well as actin to control for purity of the surface fractions. Quantification for α3- (D) and β1- (E) subunit-composed integrins from n = 3 independent experiments. (F) – (H) LecB was applied basolaterally to MDCK cells for the indicated time periods followed by basolateral application of activation-specific anti-β1-integrin antibodies to live cells. After fixation, the signal from bound anti-β1-integrin antibodies in randomly chosen regions of interest was measured and normalized to the cell number in the regions (n = 5 for one experiment). The graphs show the mean value from n = 3 experiments with the activating anti-β1-integrin antibody 9EG7 (F), and the inhibitory anti-β1-integrin antibodies mAB 13 (G) and AIIB2 (H).

In the surface biotinylation experiments we were not able to distinguish between active and inactive β1-integrins. Thus, we devised an alternative strategy in which we applied activation-specific β1-integrin antibodies to the basolateral surface of live cells. This approach revealed that LecB internalizes active and inactive β1-integrins at similar kinetics (Figs. 3F – 3H and Fig. S3F), which indicates that the activation status of β1-integrins does not play an important role in LecB-mediated integrin internalization. Taken together, LecB binds integrins, including α3β1-integrin, and causes their internalization and degradation regardless of their activation status and bound basement membrane ligands.

### 2.4 Membrane invagination by LecB and LecB-mediated crosslinking of fucosylated lipids with β1-integrin can explain LecB-triggered integrin internalization

Endogenous lectins, like galectin-3, were previously shown to be able to mediate integrin internalization (39, 40). The proposed mechanism for galectin-3-mediated integrin internalization is that galectin-3 is able to cause plasma membrane invaginations by binding to glycolipids and also drags integrins into the invaginated membrane regions by functioning as a crosslinker between glycolipids and integrins. Since LecB is a tetramer with four opposing fucose binding sites (41), which represents an ideal geometry for a potential crosslinker, we investigated if a galectin-like mechanism could explain LecB-mediated β1-integrin internalization.

In a first step we examined if binding of LecB to fucosylated glycosphingolipids is sufficient to induce membrane invaginations in giant unilamellar vesicles (GUVs) (42). Indeed, GUVs containing glycosphingolipids that bear the fucosylated Lewis a antigen (Fig. 4A) or other fucosylated glycosphingolipids (Fig. S3) showed invaginations immediately after LecB application, whereas control GUVs with the non-fucosylated glycosphingolipid lactotetraosylceramide (Lc4cer) did not (Fig. 4A). To investigate the relevance of this effect, we carried out experiments with energy-depleted cells, because under these conditions cellular machineries cannot pinch off vesicles, which previously led to well-visible membrane invaginations when other lipid-binding lectins like Shiga toxin were applied (42, 43). Indeed, LecB was able to induce plasma membrane invaginations in energy-depleted MDCK cells (Fig. 4B). Importantly, β1-integrin co-localized with fluorescently labeled LecB at invaginations (Fig. 4B, magnification), thus implicating that LecB can recruit integrins to invaginations. Furthermore, we observed that basolateral LecB application led to marked clustering of endogenous galectin-3 (Fig. S4), which could suggest that LecB outcompetes galectin-3-integrin interaction.

**Figure 4:**
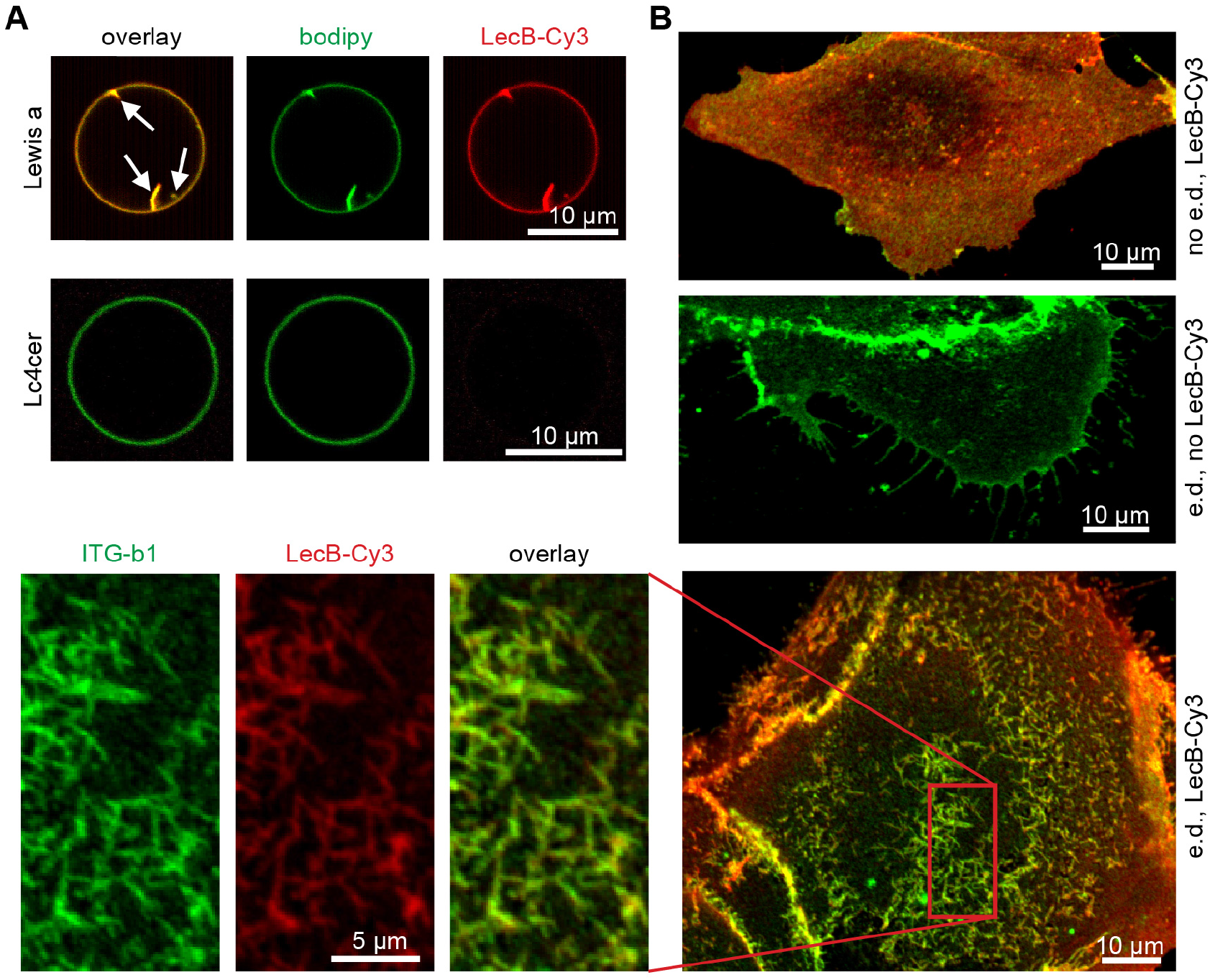
Mechanism of LecB-mediated integrin internalization via crosslinking glycosphingolipids and integrins. (A) LecB-Cy3 (15 μg/ml, red) was applied to GUVs containing fucosylated glycosphingolipids bearing the Lewis a antigen (Lewis a) or the non-fucosylated precursor lactotetraosylceramide (Lc4cer) and BODIPY-FL-C5-HPC (bodipy; green) as a membrane marker. Confocal sections along equatorial planes of the GUVs are displayed; white arrows point to membrane invaginations caused by LecB. (B) Sub-confluent MDCK cells grown on glass cover slips were energy-depleted (e.d.) or left untreated (no e.d.). LecB-Cy3 (red) was applied to the cells for 1 h, cells were fixed, and stained for β1-integrin (green). Confocal x-y sections at the level of the cell adhesion to the glass cover slip are displayed.

In summary, LecB is able to cause membrane invaginations upon binding to fucose-bearing glycosphingolipids. Since LecB is also able to bind integrins and thus to crosslink lipids in membrane invaginations with integrins, this provides a mechanistic explanation for LecB-mediated integrin internalization.

### 2.5 LecB inhibits cell migration and epithelial wound healing

As an opportunistic pathogen *P. aeruginosa* mainly relies on, and exploits, extrinsic circumstances – like a wound – to gain access to the basolateral side of epithelia. In addition, integrin blocking, e.g. through antibodies, has been previously shown to inhibit cell migration in wound healing assays (44). These considerations motivated us to investigate the effect of LecB on epithelial wound healing. Indeed, presence of LecB strongly inhibited collective cell migration and wound healing in MDCK monolayers (Fig. 5A). Importantly, this effect was blockable with L-fucose, demonstrating that LecB needs to bind to host cells to cause migration defects. Moreover, we established that the blockage of wound healing by LecB occurred in a dose-dependent manner, with concentrations larger than 50 μg/ml completely blocking cell migration (Figs. 5B and 5C), whereas another fucose-binding lectin, UEA-I (50 μg/ml), did not induce suppression of wound healing (Fig. S5A). The inhibitory effect of LecB on wound healing was – as other LecB-mediated effects before – reversible by washing out LecB (Fig. S5B).

**Figure 5:**
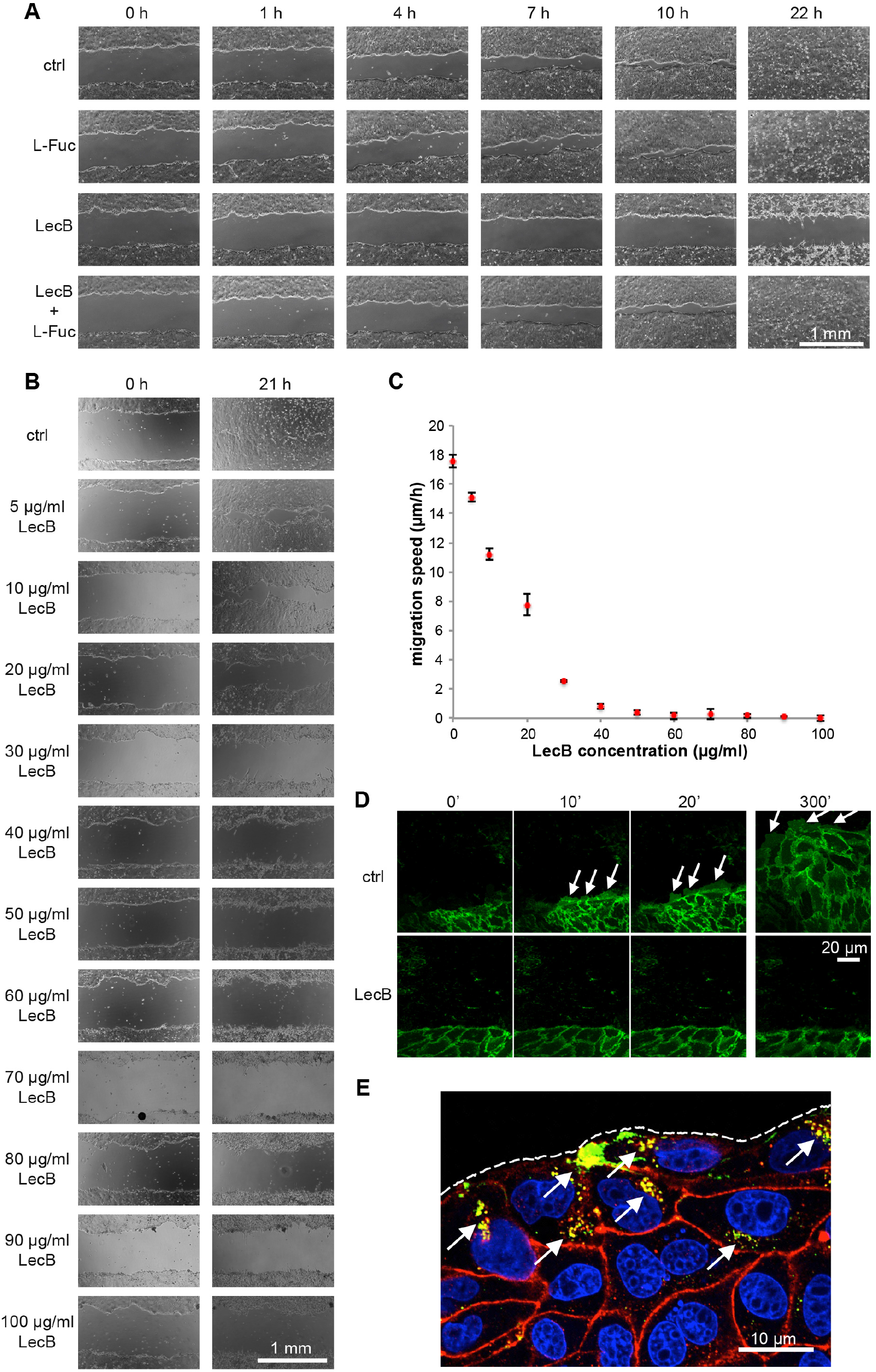
LecB inhibits epithelial wound healing. (A) – (C) Polarized monolayers of MDCK cells grown in 12 well plates were wounded with a pipet tip and imaged with a wide field microscope at the indicated time points to observe wound closure. In (A) cells were treated with LecB and/or L-fucose (43 mM) to block LecB, whereas in (B) increasing concentrations of LecB were used. The quantification of the migration speeds of the wound edges from the latter experiment (C) shows that concentrations larger than 50 μg/ml LecB completely inhibit wound healing; n = 3. (D) Polarized monolayers of MDCK cells stably expressing the plasma membrane marker ML-GFP (green) grown on chambered cover glasses were wounded and left untreated (ctrl) or treated with LecB followed by live imaging of the wound edge by confocal microscopy. Lamellipodia are indicated with white arrows. (E) Polarized MDCK monolayers grown on chambered cover glasses were wounded and treated with LecB-Alexa488 (green) for 3 h. Cells were fixed and stained for β1-integrin (red). A x-y confocal section at half height of the cells is shown. White arrows point to internalized β1-integrins co-localizing with LecB-Alexa488; the dashed line outlines the wound edge.

To explain the abrogation of cell migration by LecB we carried out live cell imaging experiments with MDCK cells stably expressing the plasma membrane marker ML-GFP (45). ML-GFP allowed visualizing lamellipodia formed by migrating MDCK cells (Fig. 5D, ctrl, white arrows). Interestingly, when cells were treated with LecB right after wounding, no lamellipodia formed (Fig. 5D, LecB), whereas washout of LecB was sufficient to reinstate lamellipodia formation and cell migration (Fig. S5C). When migrating cells were treated with LecB, lamellipodia ‘froze’ and LecB strongly bound to lamellipodia (Fig. S6). It is also interesting to note that cells deeper within the monolayer, which expose only their apical membranes to LecB in this assay, did still move (Fig. S6). In subsequent experiments we stained for β1-integrins in wound healing assays. This revealed that cells at the wound edge take up large amounts of LecB and in the same cells pronounced β1-integrin internalization was evident (Fig. 5E, white arrows), which can explain why these cells are not able to migrate any more.

Taken together, LecB inhibits epithelial wound healing in a reversible manner, which is presumably caused by the fact that integrins in wound edge cells are accessible by LecB and are internalized.

### 2.6 In dependence of LecB expression, P. aeruginosa is able to crawl underneath cells

The additional and probably dominant cytotoxic effects caused by the numerous toxins produced by *P. aeruginosa* prevented us from directly quantifying an effect of LecB knockdown in *in vitro* wound healing assays with live *P. aeruginosa*. However, we observed that *P. aeruginosa* (PAO1-wt) is able to crawl underneath exposed cells (Fig. 6A). We postulated that this requires at least local loosening of potentially integrin-mediated cell-substrate adhesion. We tested this hypothesis by investigating the influence of LecB on ‘*P. aeruginosa*-crawling’. To this end, we used a LecB-deficient PAO1 strain (PAO1-dLecB), which exhibited the same growth kinetics as PAO1-wt (Fig. S7A). After overnight culture the PAO1-wt strain showed clear expression of LecB, whereas the LecB-deficient *P. aeruginosa* (PAO1-dLecB) did not (Figs. S7B and S7C). In accordance with our hypothesis, PAO1-dLecB was visibly found less underneath cells (Fig. 6A). To substantiate the experimental procedure, we established that increasing the multiplicity of infection (MOI) (Figs. 6B and 6C) and increasing the duration of incubation (Figs. 6B and 6D) also increased the number of bacteria crawling underneath cells. Importantly, for all investigated conditions, the number of underneath-crawling bacteria per cell was lower for the PAO1-dLecB strain in comparison to the PAO1-wt strain.

**Figure 6:**
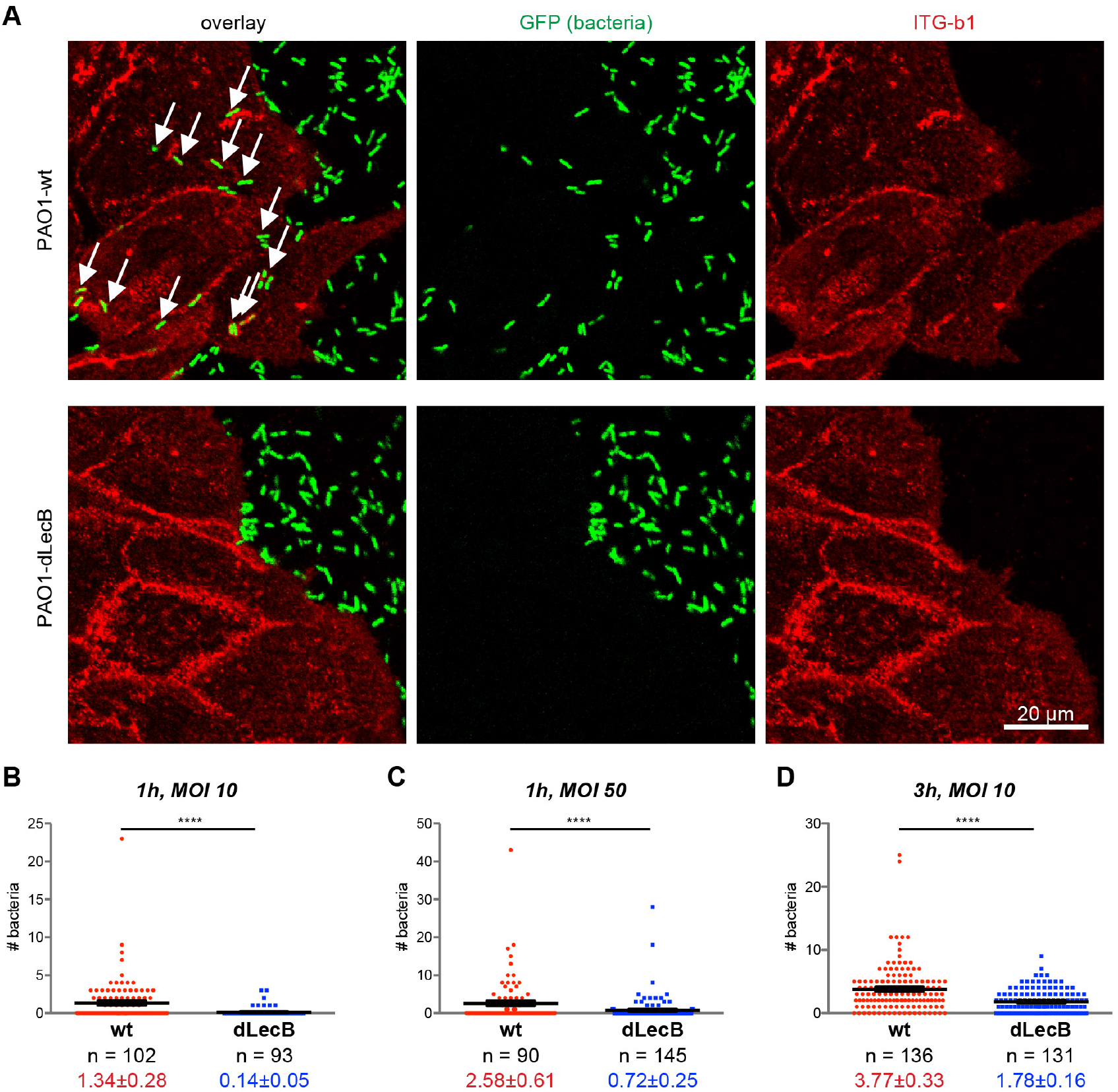
LecB promotes crawling of *P. aeruginosa* underneath cells. (A) Sparsely seeded MDCK cells were incubated with GFP-tagged PAO1-wt or LecB-deficient PAO1-dLecB (green) at an MOI of 50 for 1 h. After fixation, β1-integrins were stained in red. For each condition, a confocal x-y section at the level of cell adhesion to the substrate, which was taken from a complete three-dimensional confocal stack is displayed. White arrows indicate bacteria underneath cells. (B) – (D) After carrying out an experiment as described in (A) but with indicated MOIs and incubation periods, the numbers of bacteria underneath cells were determined per cell. Each data point represents an individual cell and the black marks indicate the mean and the SEM. For evaluating statistical significance, Mann-Whitney testing using GraphPad Prism 5 was applied, **** denotes p < 0.0001.

In summary, our study reveals a novel mechanism in which LecB, through impairing cell-to-basement membrane attachment, allows for the creation of microniches enabling sub-epithelial colonization by *P. aeruginosa*.

## 3 Discussion

### 3.1 Integrins are major receptors of LecB

Lectins bind to carbohydrates and can therefore have multiple receptors that express the appropriate glycosylation. Hence it is interesting that our experiments suggest a favored binding of LecB to integrins. LecB binding to β1-integrin appeared to be strong, because LecB was able to recover 75% of total β1-integrin from cells when it was basolaterally applied and had only access to β1-integrin at the cell surface.. As expected, LecB binding to β1-integrin solely occurred through carbohydrates, since removal of β1-integrin glycosylation abolished binding. Furthermore, mass spectrometry analysis of all basolateral LecB receptors revealed that virtually all integrins expressed by MDCK cells were among the top hits, including many integrin-associated proteins that were presumably recovered through co-precipitation with integrins.

This introduces LecB as new member to the list of bacterial molecules that bind integrins.

Another fucose binding lectin, UEA-I, was not able to mount any of the cellular effects that LecB caused. This shows that only binding to fucose is not sufficient and suggests that LecB has additional features that bring about its specific capabilities. First, LecB and UEA-I prefer different fucose linkages in oligosaccharides (UEA-I binds α_1,2_-linked fucose but only weakly to α_1,3_- and α_1,6_-linked fucose (46, 47); LecB shows a slight preference for α_1,4_-linked fucose, but also binds to α_1,2_- and α_1,3_-linked fucose (48–50)). In addition, LecB is a tetramer with four fucose binding sites, but UEA-1 is a dimer offering only two binding sites (35).

It is also worth to note that the sequence of LecB varies slightly between different strains of *P. aeruginosa* (51, 52). Since ligand binding among LecB variants is conserved, we expect that these LecB variants are utilized in similar ways as we reported here for LecB in the *P. aeruginosa* strain PAO1.

### 3.2 Mechanism of LecB-mediated integrin internalization

Since LecB recognizes only the carbohydrate moieties of integrins, it is able to manipulate integrins via unique mechanisms. The structure of LecB with four opposing binding sites is ideal to crosslink receptors. This has been demonstrated before *in vitro* by showing that LecB is able to crosslink GUVs that contain LecB receptors (53).

We hypothesize that the crosslinking capacity of LecB is key for the observed rapid internalization of integrins. In addition to integrins, LecB was also able to recognize fucose-bearing glycosphingolipids and to cause membrane invaginations without the need of additional energy input, which replicates effects caused by other glycosphingolipid-binding lectins like StxB (42), CtxB (54), or RSL (55). In addition, β1-integrin was also found on LecB-generated membrane invaginations on energy-depleted cells. This suggests that LecB causes on the one hand membrane invaginations and on the other hand is able to recruit integrins to these invaginations, thus constituting a potent endocytic mechanism. A similar mechanism was suggested for the host cell-endogenous protein galectin-3 (40). This means that bacteria have evolved molecules like LecB that can hijack this endogenous uptake route. Importantly, LecB-mediated lipid-integrin-crosslinking for internalization represents a mechanism that can explain the observed integrin internalization that occurred regardless of integrin activation status and carried also basement membrane ligands like laminins with it.

### 3.3 LecB-triggered inhibition of wound healing

LecB binding to basolateral cell surfaces caused cellular effects that could be causally linked to integrin internalization. In fully polarized epithelial cells, binding of LecB to basolateral cell surfaces, but not to apical cell surfaces, which do not contain significant amounts of integrins, led to loss of apico-basal polarity. This indicates the need of integrin internalization for dissolution of polarity. In addition, loss of polarity was reversible after washout of LecB and coincided well with the return of β1-integrin to the basolateral plasma membrane.

Our data also suggest that LecB-mediated integrin internalization is responsible for the observed block in cell migration in epithelial wound healing assays. First, integrins were preferentially internalized in wound edge cells and LecB prominently bound to lamellipodia in edge cells. This makes sense, since edge cells offer more cellular surface area for LecB binding and expose their integrins in contrast to cells deeper within the intact monolayer, which display only their apical surfaces to LecB. Second, edge cells rapidly and efficiently stopped moving upon LecB treatment, whereas other cells deeper within the monolayer preserved their capacity to move within the monolayer.

Diminished healing of *P. aeruginosa*-infected wounds was reported previously (10). However, *P. aeruginosa* possesses multiple mechanisms to manipulate and to intoxicate host cells. We therefore anticipate that, in order to inhibit wound healing, *P. aeruginosa* will apply different combinations of its arsenal, including LecB-mediated inhibition of cell migration, in dependence of the host tissue (25).

### 3.4 LecB has a role in enabling bacteria to crawl underneath host cells

Our experiments revealed a novel feature of *P. aeruginosa* behavior: We observed that bacteria frequently crawled underneath host cells. For this, at least local loosening of cell-substrate adhesion is required, which could be achieved by LecB-mediated integrin internalization. We therefore investigated the contribution of LecB to underneath-cell-crawling of the *P. aeruginosa* strain PAO1. Indeed, knocking out LecB significantly decreased crawling events. Based on our results, we suggest that *P. aeruginosa* uses LecB to manipulate integrin-basement membrane interaction to proceed along the interface between epithelial cells and the basement membrane.

In summary, our work brings integrins into focus as target of *P. aeruginosa* and provides additional rationales for the ongoing efforts to develop LecB inhibitors as additional treatment strategy to antibiotics (26–31).

## 4 Materials and Methods

### 4.1 Antibodies, plasmids and reagents

Used antibodies are listed in Table S2. The plasmid pPH-Akt-GFP encoding PH-Akt - GFP was a gift from Tamas Balla (Addgene plasmid # 51465). The plasmid encoding GFP tagged with a glycosylphosphatidylinositol (GPI) anchor (pGPI-GFP) was described before (56). The plasmid encoding for GFP tagged with a Lyn-derived myristoylation motif (pML-GFP) was a gift from Christian Wunder (Curie Institute, Paris, France).

Recombinant LecB was produced in *Escherichia coli* BL21(DE3) cells and purified with affinity columns as previously described (41). LecB and fluorophore-conjugated LecB were used at a concentration of 50 μg/ml (4.3 μM) unless stated otherwise. UEA-I and UEA-I-FITC were from Vector Labs. Cycloheximide, and L-fucose (6-Deoxy-L-galactose) were from Sigma Aldrich.

### 4.2 Mammalian cell culture, creation of stable cell lines

MDCK strain II cells were cultured in Dulbecco’s Modified Eagle’s Medium (DMEM) supplemented with 5% fetal calf serum (FCS) at 37 °C and 5% CO_2_. Unless stated otherwise 3×10^5^ MDCK cells were seeded per transwell filter (0.4 μm pore size, polycarbonate membrane, #3401 from Corning) and cultured for 4 d before experiments. For the creation of stable MDCK cell lines plasmids encoding the proteins of interest and G418 resistance (pPH-Akt-GFP, pML-GFP, and pGPI-GFP) were transfected into cells with Lipofectamine 2000 (Thermo Fisher). After allowing the cells to express the proteins overnight, they were trypsinized and plated sparsely in medium containing 1 mg/ml G418. After single colonies had formed, GFP-positive colonies were extracted with cloning rings. At least 6 colonies were extracted for each cell line, grown on transwell filters for 4 d, fixed and stained against the basolateral marker protein β-catenin and the tight junction marker protein ZO-1 to assay their polarized morphology. Based on these results we chose one colony for each cell line for further experiments. TEER measurements were carried out using an EVOM2 equipped with chopstick electrodes (World Precision Instruments).

### 4.3 Immunofluorescence

Cells were washed two times with phosphate-buffered saline without Ca^2+^ and Mg^2+^ (PBS), and then fixed with 4% (w/v) formaldehyde (FA) for 15 min at room temperature. Samples were treated with 50 mM ammonium chloride for 5 min to quench FA and then permeabilized with a SAPO medium (PBS supplemented with 0.2% (w/v) bovine serum albumin and 0.02% (w/v) saponin) for 30 min. Primary antibodies were diluted in SAPO medium and applied on the samples for 60 min at room temperature. After three washes with PBS, secondary dye-labeled antibodies, and, if required, DAPI and dye-labeled phalloidin, were diluted in SAPO medium and applied to the cells for 30 min at room temperature (details for the used antibodies are listed in Table S2). After 5 washes with PBS cells were mounted for microscopy. Since α3-integrin antibodies did not work in FA-fixed cells, methanol fixation was applied in this case. Briefly, cells were incubated with pre-cooled methanol for 15 min at −20 °C. After washing with PBS, cells were permeabilized with 0.05% (v/v) Triton X-100 for 10 min at room temperature and blocked with 3% (w/v) bovine serum albumin (BSA) for 1 h at room temperature. Staining with primary and secondary antibodies was then carried out as described before, but with a 3% (w/v) BSA solution.

### 4.4 Microscopic imaging of fixed cells and live cell experiments

For microscopic imaging an A1R confocal microscope (Nikon) equipped with a 60x oil immersion objective (NA = 1.49) and laser lines at 405 nm, 488 nm, 561 nm, and 641 nm was utilized. Image acquisition and analysis was performed with NIS-Elements 4.10.04 (Nikon).

Live cell experiments were carried out at 37°C and MDCK cells grown as polarized monolayers for 3 d on Lab-Tek II chambered cover glasses (8 well, 1.5 borosilicate glass) were used. The medium was changed to recording medium (Hank’s balanced salt solution (HBSS) supplemented with 1% FCS, 4.5 g/L glucose, and 20 mM HEPES).

### 4.5 Wound healing assays

MDCK cells were seeded on 12 well plates or, for live cell microscopy of cell migration, on 8 well Lab-Tek II chambered cover glasses and allowed to form confluent monolayers for 3 d. Then cells were scratched with a 200 μl-pipet tip to inflict a wound. On 12 well plates, marker lines were drawn on the bottom to ensure that always the same position of the wound was imaged.

### 4.6 Western blot analysis

Cells were washed twice with PBS and lysed in RIPA buffer (20 mM Tris (pH 8), 0.1% (w/v) SDS, 10% (v/v) glycerol, 13.7 mM NaCl, 2mM EDTA, and 0.5% (w/v) sodium deoxycholate in water), supplemented with protease inhibitors (0.8 μM aprotinin, 11 μM leupeptin, 200 μM pefabloc) and phosphatase inhibitor (1 mM sodium orthovanadate). Protein concentrations were analyzed using a BCA assay kit (Pierce). Equal amounts of protein per lysate were separated by SDS-PAGE and transferred to a nitrocellulose membrane. The membrane was blocked with tris-buffered saline (TBS) supplemented with 0.1% (v/v) Tween 20 and 3% (w/v) BSA for one hour and incubated with primary and HRP-linked secondary antibodies diluted in the blocking solution. In some cases, TBS supplemented with 0.1% (v/v) Tween 20 and 5% (w/v) milk powder was used (details for the used antibodies and conditions are listed in Table S2). Detection was performed by a chemiluminescence reaction using the Fusion-FX7 Advance imaging system (Peqlab Biotechnologie GmbH).

### 4.7 Energy depletion

Energy depletion was carried out as described before (42). Briefly, MDCK cells were washed 2 times with PBS supplemented with 100 mg/l CaCl_2_ and 100 mg/l MgCl_2_-6H_2_O (PBS++) and then treated with PBS++ supplemented with 10 mM 2-deoxy-D-glucose and 10mM NaN_3_ for 30 min at 37 °C.

### 4.8 Bacteria culture and crawling experiments

GFP-tagged *P. aeruginosa* PAO1 wild type (PAO1-wt) and LecB-deficient (PAO1-dLecB) strains were provided by S. de Bentzmann (CNRS - Aix Marseille University, France). The generation of LecB-deficient PAO1 is described in (23) and GFP-tagging was carried out according to the procedure described in (57). For experiments, bacteria were cultured overnight (approximately 16 h) in LB-Miller medium containing 60 μg/ml gentamicin in a shaker (Thriller, Peqlab) at 37 °C and 650 rpm. The bacteria reached an OD measured at 600 nm of approximately 5. Using these growth conditions, PAO1-wt and PAO1-dLecB strains showed comparable growth kinetics (Fig. S7A) and harvested PAO1-wt efficiently expressed LecB, whereas PAO1-dLecB did not (Figs. S7B and S7C).

For crawling experiments MDCK cells were sparsely seeded on 8 well Lab-Tek II chambered cover glasses and cultured for 1 d, so that clusters with maximally 10 cells in diameter formed to ensure that all cells were exposed to *P. aeruginosa* similar as at a wound edge. Then, PAO1-wt or PAO-dLecB were applied for the indicated MOI and duration. Bacteria crawling under cells were counted manually per cell from confocal image stacks of whole cells to ensure that only bacteria located directly underneath cells at the level of the glass cover slip were counted.

### 4.9 qPCR

PAO1-wt and PAO1-dLecB were cultured overnight as described before. RNA was extracted using TRI reagent (Sigma Aldrich). After DNase digest, 100 ng RNA was transcribed into cDNA using a first strand cDNA synthesis kit (Thermo Fisher). Then qPCR was performed on a CFX384 qPCR cycler (Bio-Rad) using a SYBR Select master mix (Thermo Fisher) and the following primers: for LecB: forward: 3’-AAGGAGTGTTCACCCTTCCC-5’, reverse: 3’-GATGACGGCGTTATTGGTGC-5’; for rpoD as reference: forward: 3’-GGGATACCTGACTTACGCGG-5’, reverse: 3’-GGGGCTGTCTCGAATACGTT-5’.

### 4.10 Labeling of lectins

LecB was labeled with fluorescent dyes bearing NHS esters as reactive groups (Cy3 mono-reactive NHS ester (GE Healthcare), Cy5 mono-reactive NHS ester (GE Healthcare), Alexa488 NHS ester (Thermo Fisher)) or with biotin using NHS-PEG12-biotin (Thermo Fisher) according to the instructions of the manufacturers and purified using PD-10 desalting columns (GE Healthcare).

### 4.11 Cell surface biotinylation and immunoprecipitation

For cell surface biotinylation, all following steps were carried out in a cold room (4 °C). Sulfo-NHS-biotin (Thermo Fisher) was freshly diluted in PBS++ (concentration 0.3 mg/ml) and applied to the apical or basolateral plasma membrane of transwell filter grown MDCK cells for 20 min. Afterwards, the reaction was quenched for 20 min with PBS++ supplemented with 50 mM ammonium chloride. Cells were lysed with RIPA buffer and biotinylated proteins were precipitated with streptavidin-agarose beads (Thermo Fisher). Elution was carried out with Laemmli buffer (2% (w/v) SDS, 10% (v/v) glycerol, 60 mM Tris-Cl (pH 6.8) in water) and boiling at 98 °C for 5 min.

For β1-integrin IP, MDCK cells were grown to confluence in 10 cm dishes and lysed in IP lysis buffer (50 mM Tris-HCl (pH 7.5), 150 mM sodium chloride, 1% (v/v) IGEPAL CA-630, 0.5% (w/v) sodium deoxycholate in water). The lysates were pre-cleared with protein A-agarose beads (Roche) for 3 h and then incubated with anti-β1-integrin antibodies (MAB2000 from Millipore) for 1 h. After adding of protein A-agarose beads overnight, beads were washed three times with IP-lysis buffer, and β1-integrin was eluted with Laemmli buffer and boiling at 98 °C for 5 min.

### 4.12 Mass spectrometry-based identification of LecB interaction partners

MDCK cells were cultured in SILAC media for 9 passages and then seeded on transwell filters and allowed to polarize for 4 d. For the first sample, biotinylated LecB was applied to the apical side of light-SILAC-labeled cells and on the basolateral side of medium-SILAC-labeled cells, whereas heavy-SILAC-labeled cells received no stimulation and served as control. For the second sample, the treatment conditions were permuted. After lysis with IP lysis buffer, the different SILAC lysates were combined and LecB-biotin-receptor complexes were precipitated using streptavidin agarose beads as described before. Eluted LecB-biotin-receptor complexes were then prepared for MS analysis using SDS-PAGE gel electrophoresis. Gels were cut into pieces, proteins therein digested with trypsin and resulting peptides were purified by STAGE tips.

For mass spectrometry analysis, samples were fractionated by nanoscale HPLC on a 1200 HPLC (Agilent Technologies, Waldbronn, Germany) connected online to a LTQ Orbitrap XL mass spectrometer (Thermo Fisher Scientific, Bremen, Germany). Fused silica HPLC-column tips with 75 μm inner diameter were self-packed with Reprosil– Pur 120 ODS–3 (Dr. Maisch, Ammerbuch, Germany) to a length of 20 cm. Samples were directly injected into the mass spectrometer. For details see reference #(58). The raw data files were uploaded into the MaxQuant software. Database searches were performed against a full-length dog database containing common contaminants such as keratins and enzymes used for in–gel digestion. Carbamidomethylcysteine was set as fixed modification, oxidation of methionine and protein amino–terminal acetylation was set as variable modifications. Triple SILAC was used as quantitation mode. The enzyme specificity was trypsin/P+DP with three allowed miss cleavages. The MS/MS tolerance was set to 0.5 Da and the mass precision of identified peptides after recalibration was in general less than 1 ppm. For identification and quantitation, the following settings were used: peptide and protein FDR were set to 0.01, maximum peptide posterior error probability (PEP) was set to 0.1, minimum peptide length was set to 7, minimum number peptides for identification and quantitation of proteins was set to two of which one must be unique, minimum ratio count was set to two, and only unmodified peptides and the variable modification were used for protein quantification. The “match between run” option was used with a time window of 2 min.

From the generated list of MS-identified proteins we defined those proteins as LecB interaction partners that showed more than 2-fold enrichment on a log2-scale over controls in both SILAC samples (see Table S1).

### 4.13 GUV experiments

Giant unilamellar vesicles (GUVs) were composed of DOPC, spiked with 1 mol% BODIPY-FL-C_5_-HPC, 30 mol% cholesterol, and 5 mol% of the desired glycosphingolipid species. Blood group glycosphingolipids were provided by Göran Larson (Sahlgrenska University Hospital, Gothenburg, Sweden). GUVs were grown at room temperature using the electroformation technique on indium-tin oxide (ITO)-coated slides as described previously (42, 43). Briefly, lipid mixtures were dissolved in chloroform at a final concentration of 0.5 mg/ml, and 15μl solution was spread on the conductive surface of ITO slides. After 2 h of drying under vacuum, GUVs were grown in a 290 mosM sucrose solution by applying an alternating electric field from 20 mV to 1.1 V for 3 h.

LecB-Cy3 (15 μg/ml) was incubated with GUVs at room temperature and examined under an inverted confocal fluorescence microscope (Nikon A1R) equipped with an oil immersion objective (60x, NA 1.49).

### 4.14 Statistics

If not stated otherwise data obtained from n = 3 independent experiments were used to calculate arithmetic means. Error bars represent standard error mean (SEM). Statistical significance analysis was carried out using GraphPad Prism 5.

## 5 Acknowledgements

This work was supported by a German Research Foundation grant [RO 4341/2-1], the Excellence Initiative of the German Research Foundation [EXC 294], the Ministry of Science, Research and the Arts of Baden-Württemberg [Az: 33-7532.20], and a starting grant from the European Research Council [Programme “Ideas,” ERC-2011-StG 282105]. R.T. acknowledges support from the Ministry of Science, Research and the Arts of Baden-Württemberg [Az: 7533-30-10/25/36].

## References

1. Lyczak JB, Cannon CL, Pier GB. 2002. Lung infections associated with cystic fibrosis. Clin Microbiol Rev 15:194–222.

2. Franzetti F, Grassini A, Piazza M, Degl’Innocenti M, Bandera A, Gazzola L, Marchetti G, Gori A. 2006. Nosocomial bacterial pneumonia in HIV-infected patients: risk factors for adverse outcome and implications for rational empiric antibiotic therapy. Infection 34:9–16.

3. Markou P, Apidianakis Y. 2014. Pathogenesis of intestinal Pseudomonas aeruginosa infection in patients with cancer. Front Cell Infect Microbiol 3:115.

4. Barbier F, Andremont A, Wolff M, Bouadma L. 2013. Hospital-acquired pneumonia and ventilator-associated pneumonia: recent advances in epidemiology and management. Curr Opin Pulm Med 19:216–228.

5. Azzopardi EA, Azzopardi E, Camilleri L, Villapalos J, Boyce DE, Dziewulski P, Dickson WA, Whitaker IS. 2014. Gram negative wound infection in hospitalised adult burn patients-systematic review and metanalysis. PLoS One 9:e95042.

6. Stover C, Pham X, Erwin A, Mizoguchi S, Warrener P, Hickey M, Brinkman F, Hufnagle W, Kowalik D, Lagrou M, Garber R, Goltry L, Tolentino E, Westbrock-Wadman S, Yuan Y, Brody L, Coulter S, Folger K, Kas A, Larbig K, Lim R, Smith K, Spencer D, Wong G, Wu Z, Paulsen I, Reizer J, Saier M, Hancock R, Lory S, Olson M. 2000. Complete genome sequence of Pseudomonas aeruginosa PAO1, an opportunistic pathogen. Nature 406:959–964.

7. Maillet M, Pelloux I, Forli A, Vancoetsem K, Cheong Sing JS, Marfaing S, Ducki S, Batailler P, Mallaret M-R. 2014. Nosocomial transmission of carbapenem-resistant Pseudomonas aeruginosa among burn patients. Infect Control Hosp Epidemiol 35:597–599.

8. Kinsey CB, Koirala S, Solomon B, Rosenberg J, Robinson BF, Neri A, Halpin AL, Arduino MJ, Moulton-Meissner H, Noble-Wang J, others. 2017. Pseudomonas aeruginosa Outbreak in a Neonatal Intensive Care Unit Attributed to Hospital Tap Water. Infect Control Hosp Epidemiol 38:801–808.

9. Mah T-F, Pitts B, Pellock B, Walker GC, Stewart PS, O’Toole GA. 2003. A genetic basis for Pseudomonas aeruginosa biofilm antibiotic resistance. Nature 426:306.

10. Engel J, Eran Y. 2011. Subversion of mucosal barrier polarity by Pseudomonas aeruginosa. Front Microbiol 2:1–7.

11. WHO. 2017. WHO priority pathogens list for research and development of new antibiotics.

12. Wagner S, Sommer R, Hinsberger S, Lu C, Hartmann RW, Empting M, Titz A. 2016. Novel Strategies for the Treatment of Pseudomonas aeruginosa Infections. J Med Chem 59:5929–5969.

13. Golovkine G, Faudry E, Bouillot S, Elsen S, Attrée I, Huber P. 2016. Pseudomonas aeruginosa transmigrates at epithelial cell-cell junctions, exploiting sites of cell division and senescent cell extrusion. PLoS Pathog 12:e1005377.

14. Tran CS, Eran Y, Ruch TR, Bryant DM, Datta A, Brakeman P, Kierbel A, Wittmann T, Metzger RJ, Mostov KE, Engel JN. 2014. Host cell polarity proteins participate in innate immunity to Pseudomonas aeruginosa infection. Cell Host Microbe 15:636–43.

15. Kwok T, Zabler D, Urman S, Rohde M, Hartig R, Wessler S, Misselwitz R, Berger J, Sewald N, Konig W, Backert S. 2007. Helicobacter exploits integrin for type IV secretion and kinase activation. Nature 449:862–866.

16. Stewart PL, Nemerow GR. 2007. Cell integrins: commonly used receptors for diverse viral pathogens. Trends Microbiol 15:500–507.

17. Cossart P. 1997. Host/pathogen interactions. Subversion of the mammalian cell cytoskeleton by invasive bacteria. J Clin Invest 99:2307–2311.

18. Roger P, Puchelle E, Bajolet-Laudinat O, Tournier J-M, Debordeaux C, Plotkowski M-C, Cohen JHM, Sheppard D, De Bentzmann S. 1999. Fibronectin and α5β1 integrin mediate binding of Pseudomonas aeruginosa to repairing airway epithelium. Eur Respir J 13:1301–1309.

19. Leroy-Dudal J, Gagnière H, Cossard E, Carreiras F, Di Martino P. 2004. Role of αvβ5 integrins and vitronectin in Pseudomonas aeruginosa PAK interaction with A549 respiratory cells. Microbes Infect 6:875–881.

20. Imberty A, Wimmerová M, Mitchell EP, Gilboa-Garber N. 2004. Structures of the lectins from Pseudomonas aeruginosa: Insights into the molecular basis for host glycan recognition. Microbes Infect 6:221–228.

21. Tielker D, Hacker S, Loris R, Strathmann M, Wingender J, Wilhelm S, Rosenau F, Jaeger KE. 2005. Pseudomonas aeruginosa lectin LecB is located in the outer membrane and is involved in biofilm formation. Microbiology 151:1313–1323.

22. Funken H, Bartels KM, Wilhelm S, Brocker M, Bott M, Bains M, Hancock REW, Rosenau F, Jaeger KE. 2012. Specific Association of Lectin LecB with the Surface of Pseudomonas aeruginosa: Role of Outer Membrane Protein OprF. PLoS One 7:1–8.

23. Chemani C, Imberty A, De Bentzmann S, Pierre M, Wimmerová M, Guery BP, Faure K. 2009. Role of LecA and LecB lectins in Pseudomonas aeruginosa-induced lung injury and effect of carbohydrate ligands. Infect Immun 77:2065–2075.

24. Adam EC, Mitchell BS, Schumacher DU, Grant G, Schumacher U. 1997. Pseudomonas aeruginosa II lectin stops human ciliary beating: Therapeutic implications of fucose. Am J Respir Crit Care Med 155:2102–2104.

25. Cott C, Thuenauer R, Landi A, Kühn K, Juillot S, Imberty A, Madl J, Eierhoff T, Römer W. 2016. Pseudomonas aeruginosa lectin LecB inhibits tissue repair processes by triggering β-catenin degradation. Biochim Biophys Acta (BBA)-Molecular Cell Res 1863:1106–1118.

26. Sommer R, Wagner S, Rox K, Varrot A, Hauck D, Wamhoff E-C, Schreiber J, Ryckmans T, Brunner T, Rademacher C, others. 2018. Glycomimetic, orally bioavailable LecB inhibitors block biofilm formation of Pseudomonas aeruginosa. J Am Chem Soc 140:2537–2545.

27. Sommer R, Hauck D, Varrot A, Wagner S, Audfray A, Prestel A, Möller HM, Imberty A, Titz A. 2015. Cinnamide Derivatives of d-Mannose as Inhibitors of the Bacterial Virulence Factor LecB from Pseudomonas aeruginosa. ChemistryOpen 4:756–767.

28. Johansson EM V, Crusz SA, Kolomiets E, Buts L, Kadam RU, Cacciarini M, Bartels K-M, Diggle SP, Cámara M, Williams P, others. 2008. Inhibition and dispersion of Pseudomonas aeruginosa biofilms by glycopeptide dendrimers targeting the fucose-specific lectin LecB. Chem Biol 15:1249–1257.

29. Donnier-Marechal M, Abdullayev S, Bauduin M, Pascal Y, Fu M-Q, He X-P, Gillon E, Imberty A, Kipnis E, Dessein R, Vidal S. 2018. Tetraphenylethylene-based glycoclusters with aggregation-induced emission (AIE) properties as high-affinity ligands of bacterial lectins. Org Biomol Chem 16:8804–8809.

30. Bucher KS, Babic N, Freichel T, Kovacic F, Hartmann L. 2018. Monodisperse Sequence-Controlled alpha-l-Fucosylated Glycooligomers and Their Multivalent Inhibitory Effects on LecB. Macromol Biosci e1800337.

31. Dupin L, Noel M, Bonnet S, Meyer A, Gehin T, Bastide L, Randriantsoa M, Souteyrand E, Cottin C, Vergoten G, Vasseur J-J, Morvan F, Chevolot Y, Darblade B. 2018. Screening of a library of oligosaccharides targeting lectin LecB of Pseudomonas aeruginosa and synthesis of high affinity oligoglycoclusters. Molecules 23.

32. Rodriguez-Boulan E, Kreitzer G, Müsch A. 2005. Organization of vesicular trafficking in epithelia. Nat Rev Mol Cell Biol 6:233–247.

33. Thuenauer R, Hsu YC, Carvajal-Gonzalez JM, Deborde S, Chuang J, Römer W, Sonnleitner A, Rodriguez-Boulan E, Sung C. 2014. Four-dimensional live imaging of apical biosynthetic trafficking reveals a post-Golgi sorting role of apical endosomal intermediates. Proc Natl Acad Sci U S A 111:4127–32.

34. Kierbel A, Gassama-Diagne A, Rocha C, Radoshevich L, Olson J, Mostov K, Engel J. 2007. Pseudomonas aeruginosa exploits a PIP3-dependent pathway to transform apical into basolateral membrane. J Cell Biol 177:21–27.

35. Audette GF, Vandonselaar M, Delbaere LTJ. 2000. The 2.2 Å resolution structure of the O (H) blood-group-specific lectin I from Ulex europaeus. J Mol Biol 304:423–433.

36. Myllymäki SM, Teräväinen TP, Manninen A. 2011. Two distinct integrin-mediated mechanisms contribute to apical lumen formation in Epithelial cells. PLoS One 6.

37. Greciano PG, Moyano J V, Buschmann MM, Tang J, Lu Y, Rudnicki J, Manninen A, Matlin KS. 2012. Laminin 511 partners with laminin 332 to mediate directional migration of Madin-Darby canine kidney epithelial cells. Mol Biol Cell 23:121–136.

38. Le Bivic A, Sambuy Y, Mostov K, Rodriguez-Boulan E. 1990. Vectorial targeting of an endogenous apical membrane sialoglycoprotein and uvomorulin in MDCK cells. J Cell Biol 110:1533–1539.

39. Furtak V, Hatcher F, Ochieng J. 2001. Galectin-3 mediates the endocytosis of β-1 integrins by breast carcinoma cells. Biochem Biophys Res Commun 289:845–850.

40. Lakshminarayan R, Wunder C, Becken U, Howes MT, Benzing C, Arumugam S, Sales S, Ariotti N, Chambon V, Lamaze C, others. 2014. Galectin-3 drives glycosphingolipid-dependent biogenesis of clathrin-independent carriers. Nat Cell Biol 16:592.

41. Mitchell EP, Sabin C, Šnajdrová L, Pokorná M, Perret S, Gautier C, Hofr C, Gilboa-Garber N, Koča J, Wimmerová M, Imberty A. 2005. High affinity fucose binding of Pseudomonas aeruginosa lectin PA-IIL: 1.0 Å resolution crystal structure of the complex combined with thermodynamics and computational chemistry approaches. Proteins Struct Funct Genet 58:735–746.

42. Römer W, Berland L, Chambon V, Gaus K, Windschiegl B, Tenza D, Aly MRE, Fraisier V, Florent J-C, Perrais D, Lamaze C, Raposo G, Steinem C, Sens P, Bassereau P, Johannes L. 2007. Shiga toxin induces tubular membrane invaginations for its uptake into cells. Nature 450:670–675.

43. Römer W, Pontani LL, Sorre B, Rentero C, Berland L, Chambon V, Lamaze C, Bassereau P, Sykes C, Gaus K, Johannes L. 2010. Actin Dynamics Drive Membrane Reorganization and Scission in Clathrin-Independent Endocytosis. Cell 140:540–553.

44. Wood S, Sivaramakrishnan G, Engel J, Shafikhani SH. 2011. Cell migration regulates the kinetics of cytokinesis. Cell Cycle 10:648–654.

45. Thuenauer R, Nicklaus S, Frensch M, Troendle K, Madl J, Römer W. 2018. A microfluidic biochip for locally confined stimulation of cells within an epithelial monolayer. RSC Adv 8:7839–7846.

46. Baldus SE, Thiele J, Park Y-O, Hanisch F-G, Bara J, Fischer R. 1996. Characterization of the binding specificity of Anguilla anguilla agglutinin (AAA) in comparison to Ulex europaeus agglutinin I (UEA-I). Glycoconj J 13:585–590.

47. Sughii S, Kabat E, Hh B. 1982. Further immunochemical studies on the combining sites of Lotus tetragonolobus and Ulex europaeus I and II lectins. Carbohydr Res 99:99–101.

48. Marotte K, Sabin C, Prville C, Moumé-Pymbock M, Wimmerová M, Mitchell EP, Imberty A, Roy R. 2007. X-ray structures and thermodynamics of the interaction of PA-IIL from Pseudomonas aeruginosa with disaccharide derivatives. ChemMedChem 2:1328–1338.

49. Perret S, Sabin C, Dumon C, Pokorná M, Gautier C, Galanina O, Ilia S, Bovin N, Nicaise M, Desmadril M, others. 2005. Structural basis for the interaction between human milk oligosaccharides and the bacterial lectin PA-IIL of Pseudomonas aeruginosa. Biochem J 389:325–332.

50. Mitchell E, Houles C, Sudakevitz D, Wimmerova M, Gautier C, Pérez S, Wu AM, Gilboa-Garber N, Imberty A. 2002. Structural basis for oligosaccharide-mediated adhesion of Pseudomonas aeruginosa in the lungs of cystic fibrosis patients. Nat Struct Mol Biol 9:918.

51. Boukerb AM, Decor A, Ribun S, Tabaroni R, Rousset A, Commin L, Buff S, Doleans-Jordheim A, Vidal S, Varrot A, Imberty A, Cournoyer B. 2016. Genomic Rearrangements and Functional Diversification of lecA and lecB Lectin-Coding Regions Impacting the Efficacy of Glycomimetics Directed against Pseudomonas aeruginosa. Front Microbiol 7:811.

52. Sommer R, Wagner S, Varrot A, Nycholat CM, Khaledi A, Haussler S, Paulson JC, Imberty A, Titz A. 2016. The virulence factor LecB varies in clinical isolates: consequences for ligand binding and drug discovery. Chem Sci 7:4990–5001.

53. Villringer S, Madl J, Sych T, Manner C, Imberty A, Römer W. 2018. Lectin-mediated protocell crosslinking to mimic cell-cell junctions and adhesion. Sci Rep 8:1932.

54. Ewers H, Römer W, Smith AE, Bacia K, Dmitrieff S, Chai W, Mancini R, Kartenbeck J, Chambon V, Berland L, Oppenheim A, Schwarzmann G, Feizi T, Schwille P, Sens P, Helenius A, Johannes L. 2010. GM1 structure determines SV40-induced membrane invagination and infection. Nat Cell Biol 12:11–18; sup pp 1-12.

55. Arnaud J, Tröndle K, Claudinon J, Audfray A, Varrot A, Römer W, Imberty A. 2014. Membrane deformation by neolectins with engineered glycolipid binding sites. Angew Chemie - Int Ed 53:9267–9270.

56. Thuenauer R, Juhasz K, Mayr R, Frühwirth T, Lipp A-M, Balogi Z, Sonnleitner A. 2011. A PDMS-based biochip with integrated sub-micrometre position control for TIRF microscopy of the apical cell membrane. Lab Chip 11:3064–3071.

57. Eierhoff T, Bastian B, Thuenauer R, Madl J, Audfray A, Aigal S, Juillot S, Rydell GE, Müller S, de Bentzmann S, Imberty A, Fleck C, Römer W. 2014. A lipid zipper triggers bacterial invasion. Proc Natl Acad Sci U S A 111:6–11.

58. Thriene K, Gruning BA, Bornert O, Erxleben A, Leppert J, Athanasiou I, Weber E, Kiritsi D, Nystrom A, Reinheckel T, Backofen R, Has C, Bruckner-Tuderman L, Dengjel J. 2018. Combinatorial Omics Analysis Reveals Perturbed Lysosomal Homeostasis in Collagen VII-deficient Keratinocytes. Mol Cell Proteomics 17:565–579.

