## Supplemental Material for "The *Pseudomonas aeruginosa* lectin LecB causes integrin internalization to facilitate crawling of bacteria underneath host cells"

Germany).

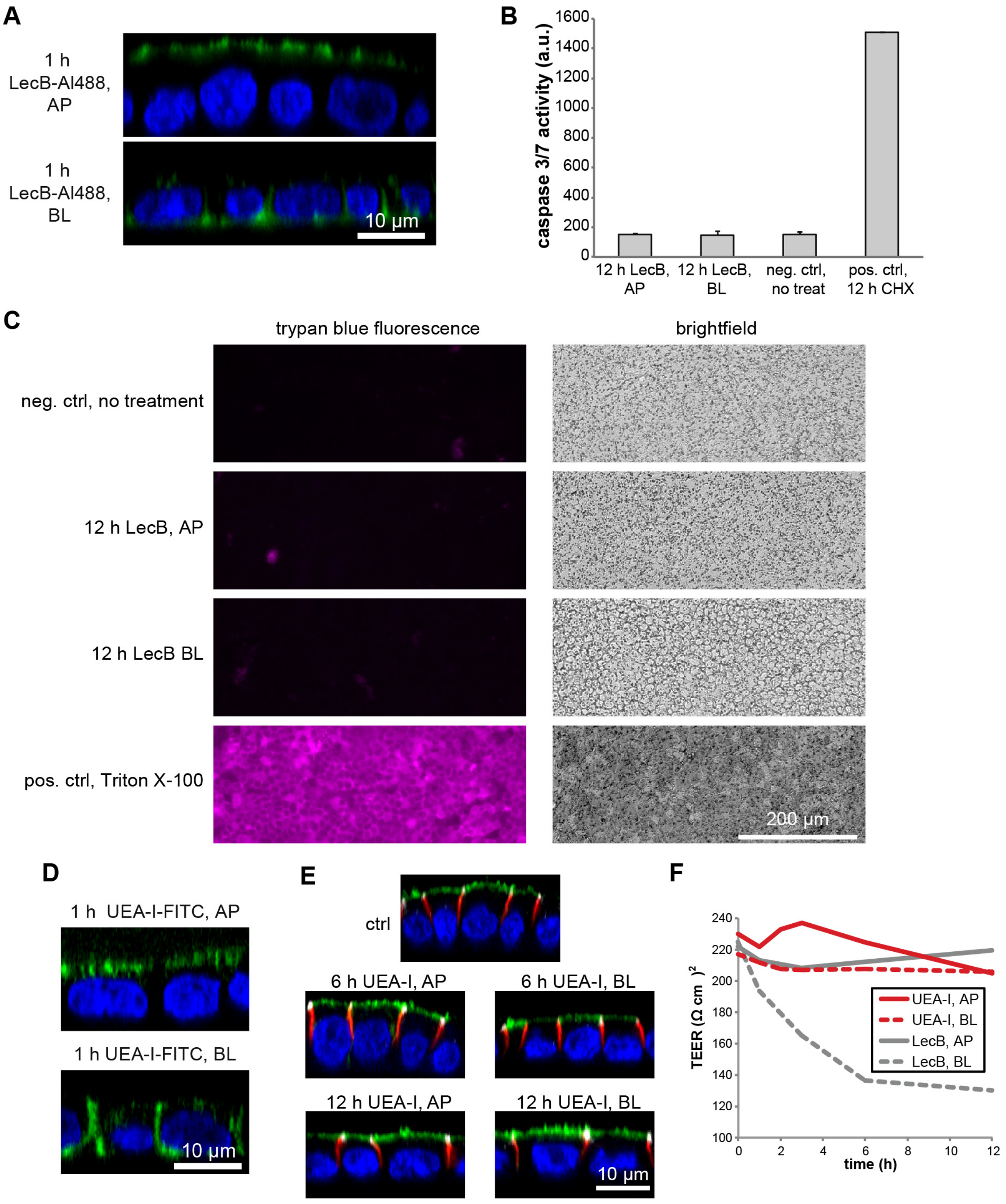

***Figure S1: Control experiments related to Fig. 1***

(A) 50 µg/ml LecB-Alexa488 (green) was applied to polarized filter-grown MDCK cells apically (AP) or basolaterally (BL). After fixation, nuclei were stained with DAPI (blue). Confocal x-z sections are displayed. (B) MDCK cells were left untreated as negative control, treated with 50 µg/ml LecB as indicated, or treated with cycloheximide (CHX; 100 µg/ml) as positive control. After cell lysis a Caspase-Glo 3/7 assay (Promega) was performed to measure the level of induced apoptosis. (C) Polarized filter-grown MDCK cells were left untreated (neg. ctrl), treated AP or BL with LecB for 12 h, or treated with 0.1% Triton X-100 for 5 min (pos. ctrl.). Afterwards cells were incubated AP and BL for 5 min with 0.4% trypan blue and then washed 2 times with PBS. Since trypan blue is fluorescent upon excitation with far-red light, its presence in cells was measured using a wide field microscope equipped with a far-red (Cy5)-cube (magenta). Cells were also imaged in brightfield mode. In these images the pores of the transwell filters are visible as small black dots. (D) 50 µg/ml UEA-I-FITC (green) was applied to polarized filter-grown MDCK cells apically (AP) or basolaterally (BL). After fixation nuclei were stained with DAPI (blue). Confocal x-z sections are displayed. (E) Polarized, filter-grown MDCK cells stably expressing the apical marker GPI-GFP (green) were left untreated (ctrl) or treated apically (AP) or basolaterally (BL) with 50 µg/ml UEA-I for the indicated time periods, fixed and stained with antibodies recognizing the basolateral marker β-catenin (red) and the tight junction marker ZO-1 (white), nuclei were stained with DAPI (blue). Representative confocal x-z sections are shown. (F) MDCK cells were treated with 50 µg/ml UEA-I as indicated and the trans-epithelial electrical resistance (TEER) was measured. As comparison the data from treating cells with LecB from Fig. 1B are shown in gray.

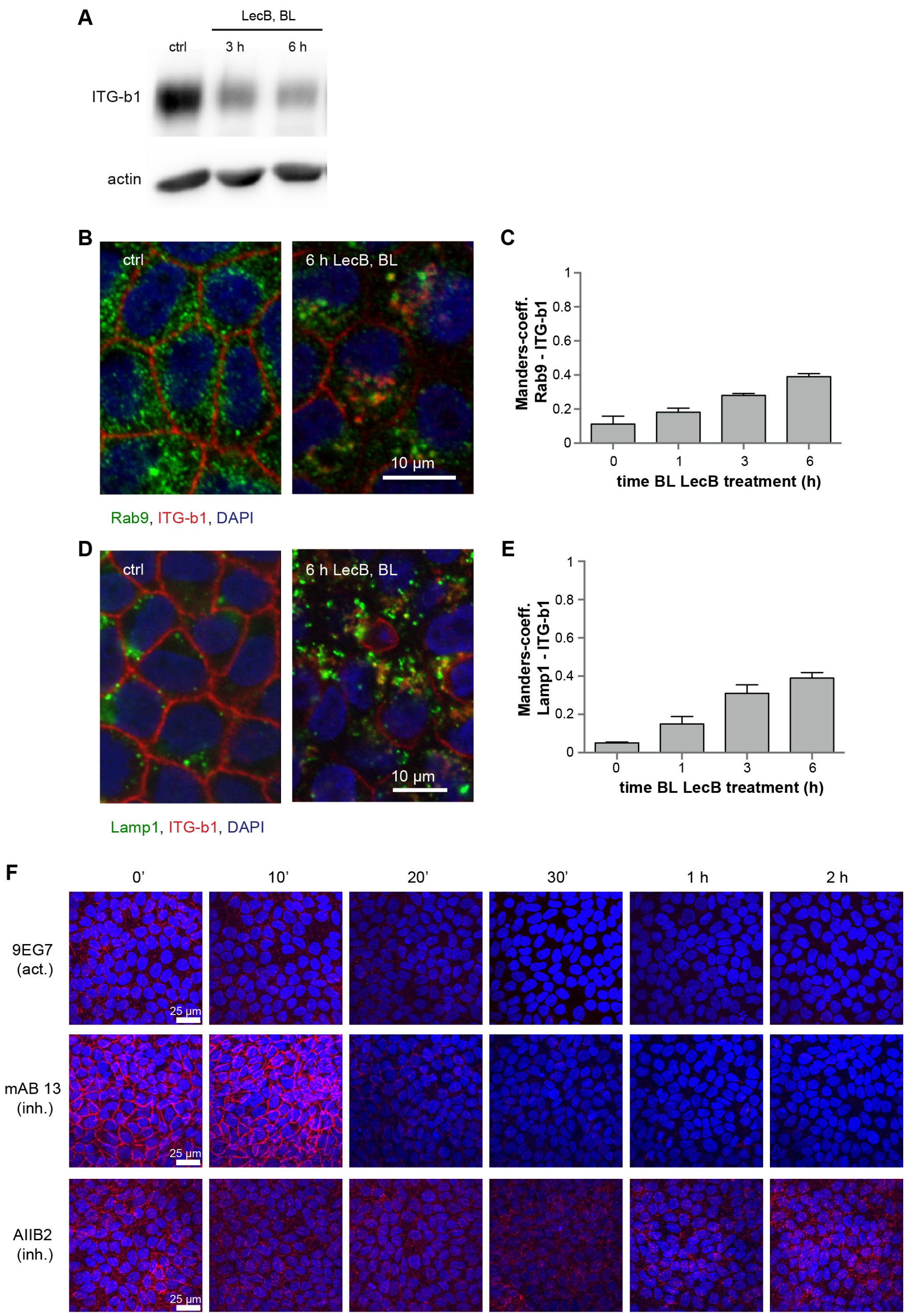

***Figure S2: Control experiments related to Fig. 3***

(A) The total amounts of β1-integrin were probed by WB in MDCK cells basolaterally treated with LecB. (B) – (C) MDCK cells were basolaterally treated with LecB as indicated, fixed, stained for Rab9 (green), β1-integrin (red), and nuclei were stained with DAPI (blue). Representative confocal sections (x-y sections) through the middle of the cells are depicted in (B) and the quantification of the Manders overlap-coefficient between Rab9 and β1-integrin from n = 3 independent experiments is depicted in (C). (D) – (E) MDCK cells were basolaterally treated with LecB as indicated, fixed, stained for Lamp1 (green), β1-integrin (red), and nuclei were stained with DAPI (blue). Representative confocal sections (x-y sections) through the middle of the cells are shown in (D) and the quantification of the Manders overlap-coefficient between Lamp1 and β1-integrin from n = 3 independent experiments is depicted in (E). (F) LecB was applied basolaterally to polarized filter-grown MDCK cells for the indicated time periods followed by basolateral application of activation-specific anti-β1-integrin antibodies (9EG7, mAB 13, AIIB2) to live cells. After fixation, the signal from bound anti-β1-integrin antibodies was measured with a confocal microscope. Representative maximum intensity projections of confocal image stacks covering full cell heights are displayed.

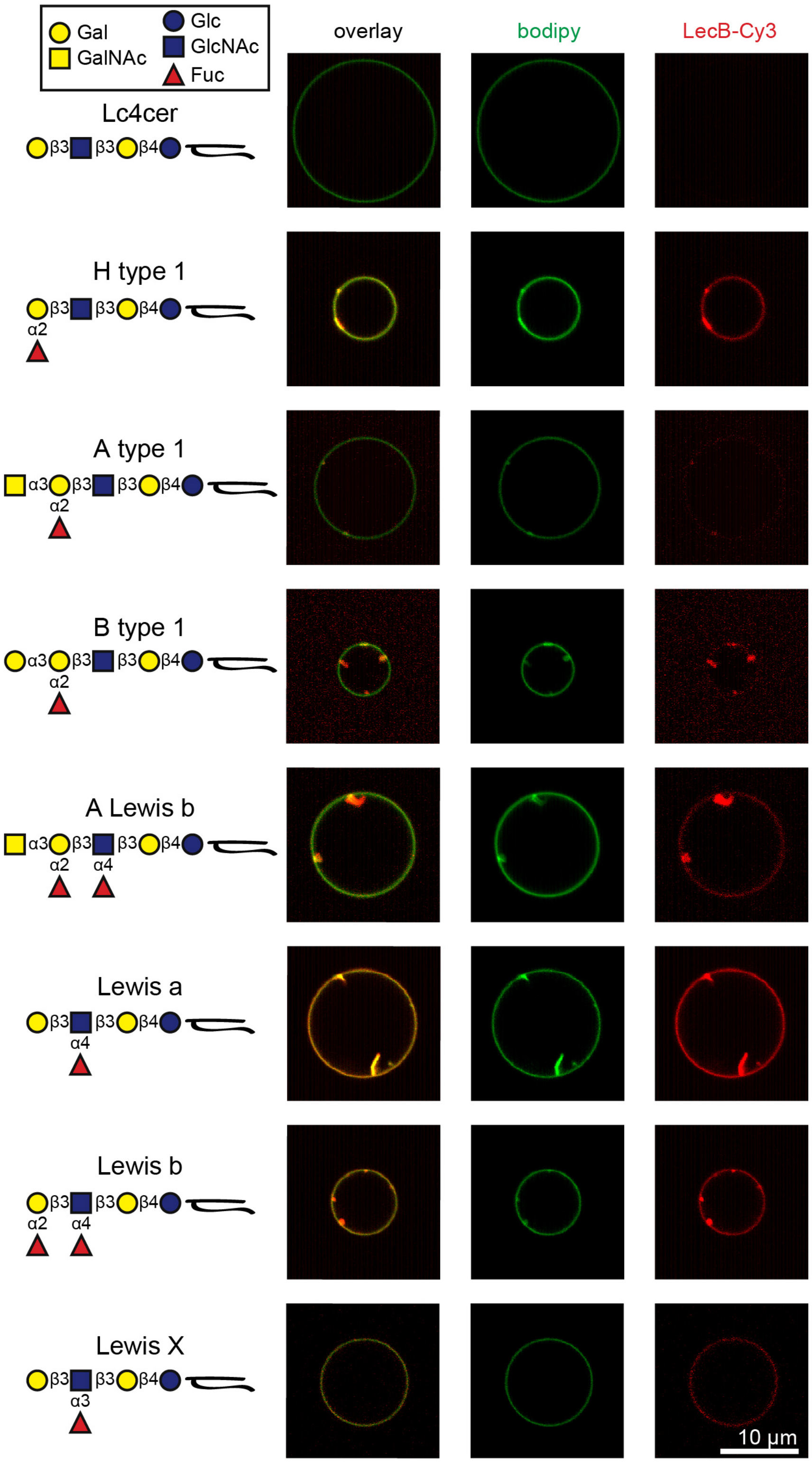

***Figure S3: Control experiments related to Fig. 4, part 1***

LecB-Cy3 (15 µg/ml, red) was applied to GUVs containing BODIPY-FL-C5-HPC (bodipy; green) as a membrane marker and glycosphingolipids bearing antigens from the type 1 series (lactotetraosylceramide (Lc4cer; non-fucosylated precursor as negative control), H type 1, A type 1, B type 1, A Lewis b, Lewis a, or Lewis b) or the Lewis X antigen from the type 2 series. Schematic structures of the glycolipids are also depicted on the left. Confocal sections along equatorial planes of representative GUVs are displayed.

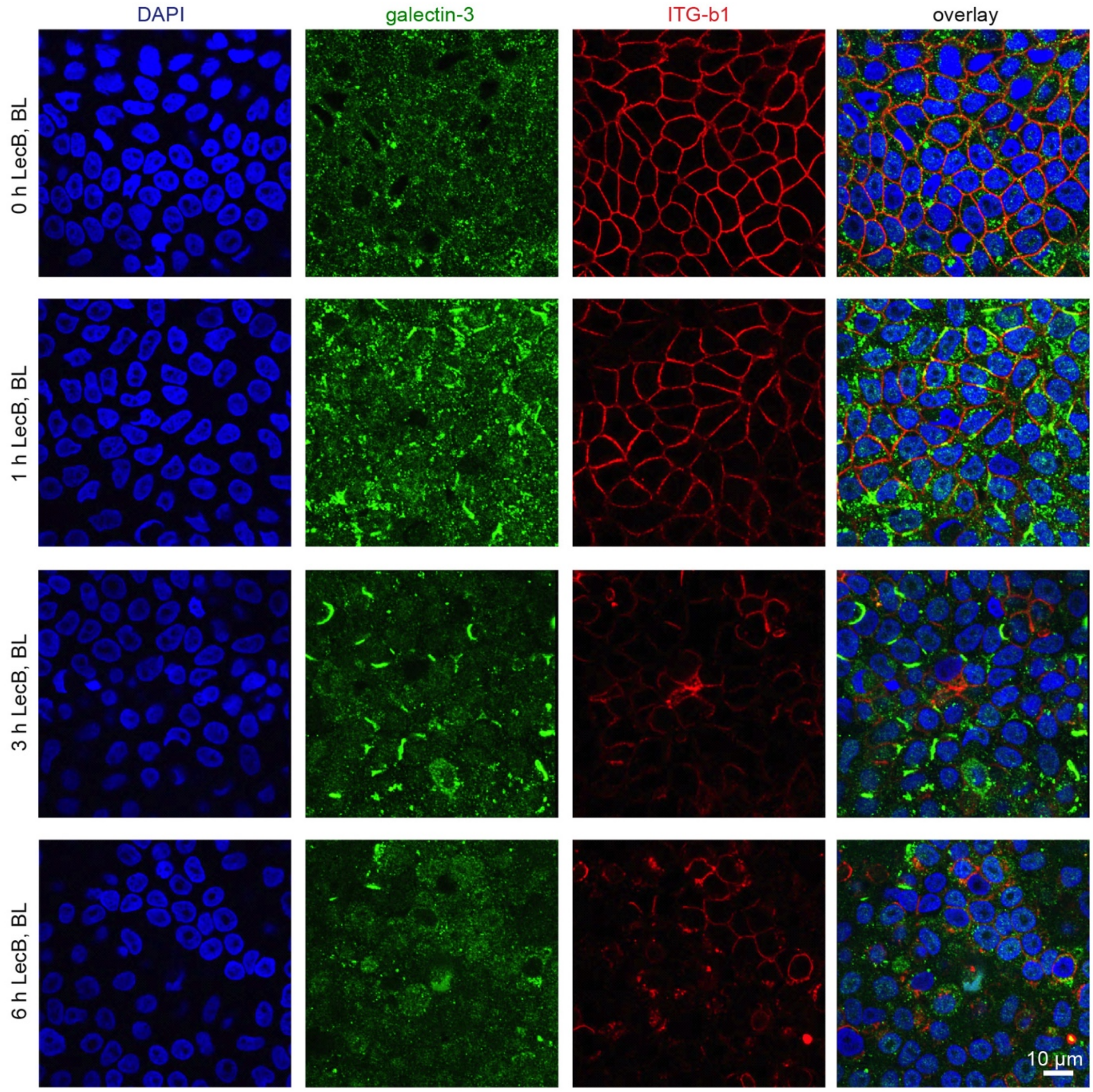

***Figure S4: Control experiments related to Fig. 4, part 2***

LecB was applied basolaterally to polarized MDCK cells grown on transwell filters. After fixation, endogenous galectin-3 (green), β1-integrin (red), and nuclei (blue) were stained. Representative confocal sections (x-y sections) from a z-level 3 µm above the transwell filter surface are displayed.

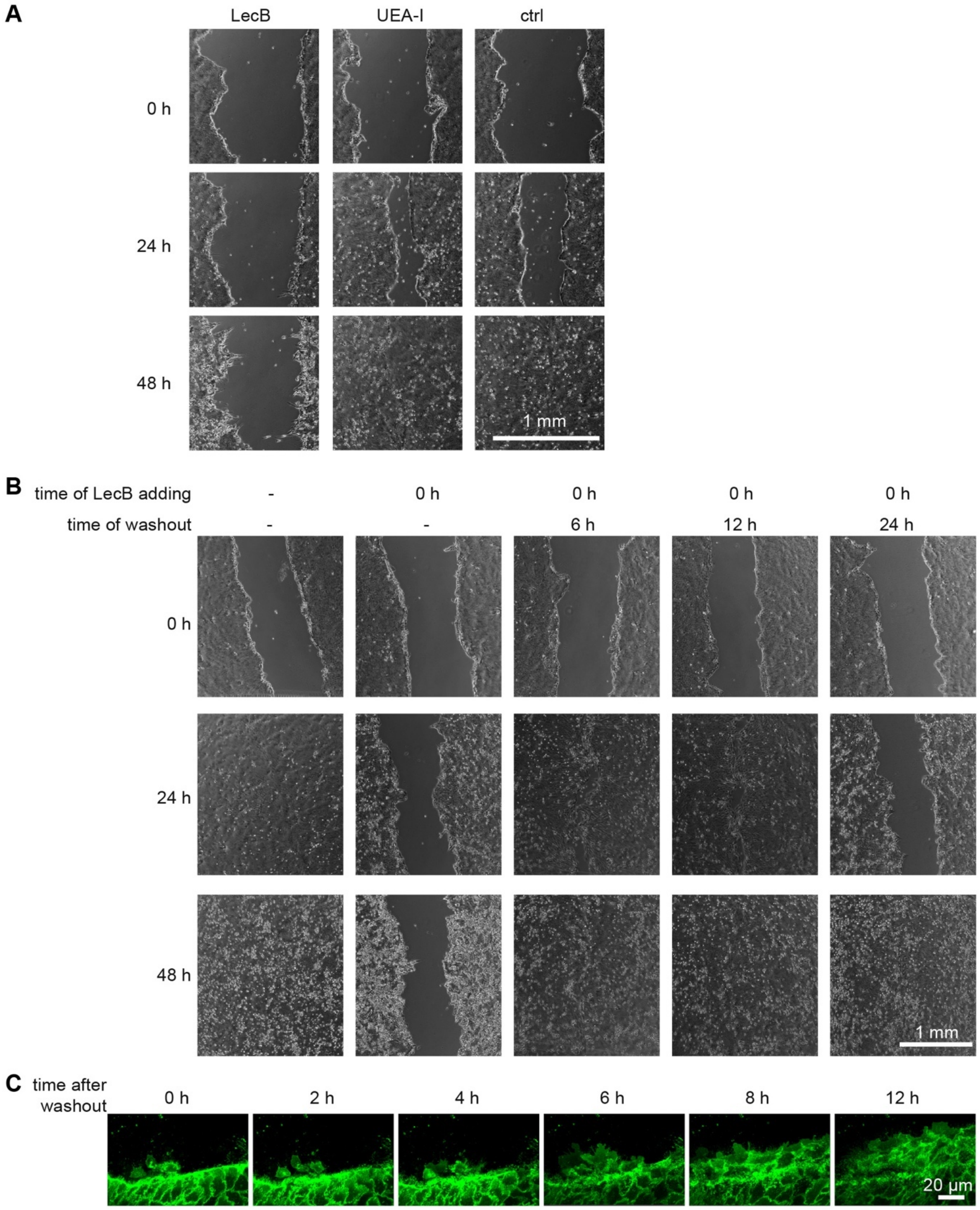

***Figure S5: Control experiments related to Fig. 5, part 1***

(A) Wound healing assays with monolayers of MDCK cells treated with 50 µg/ml UEA-I or LecB. (B) Wound healing assays with monolayers of MDCK cells in which LecB was washed out again as indicated. (C) Regeneration of cell migration at the wound edge of MDCK cells stably expressing ML-GFP (green) treated for 6 h with LecB and then washed out for the indicated times.

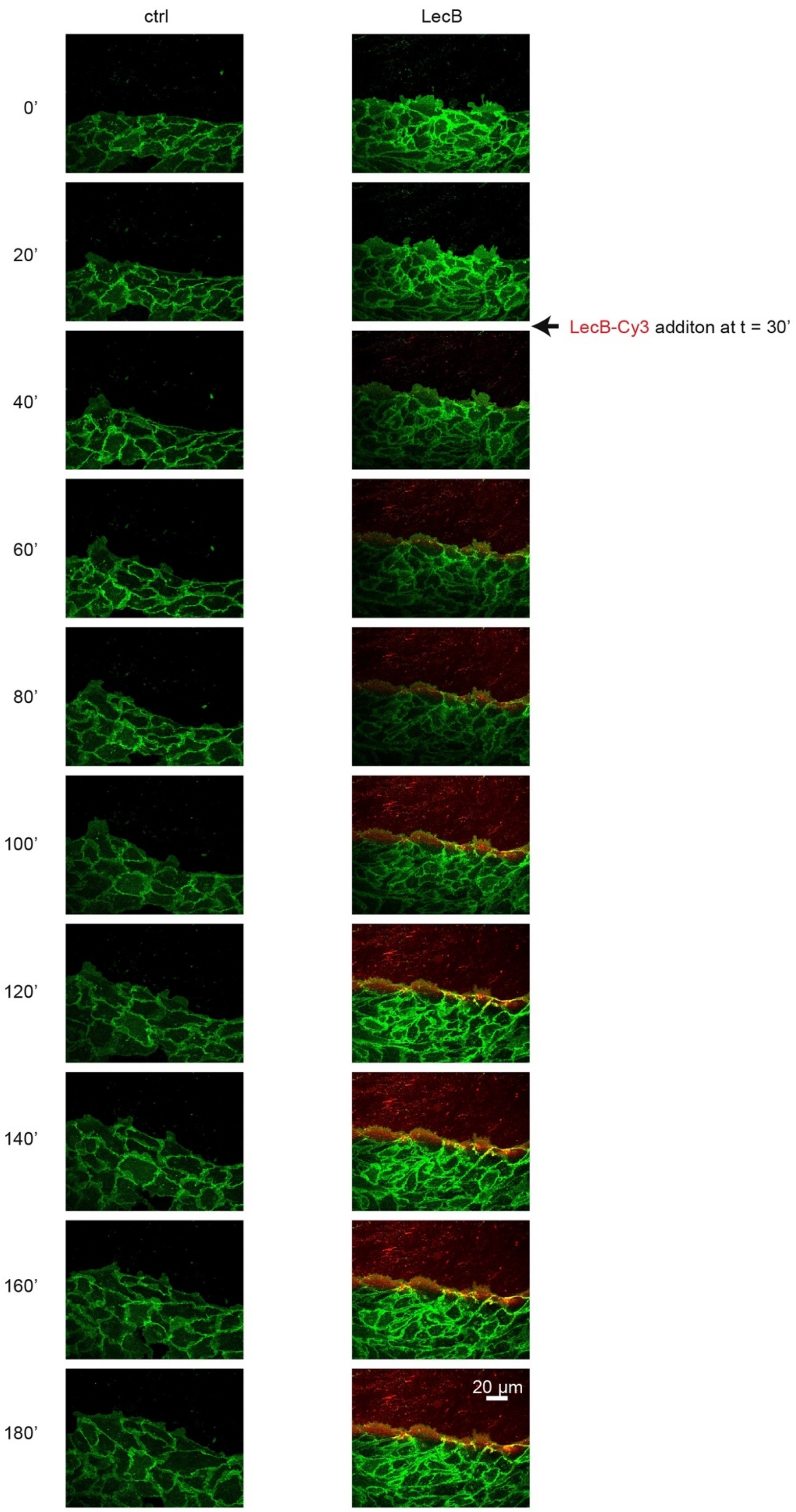

***Figure S6: Control experiments related to Fig. 5, part 2***

MDCK cells stably expressing ML-GFP (green) were wounded and observed with a confocal microscope. After 30 min, LecB-Cy3 (red) was added to one sample, whereas the other sample was left untreated (ctrl).

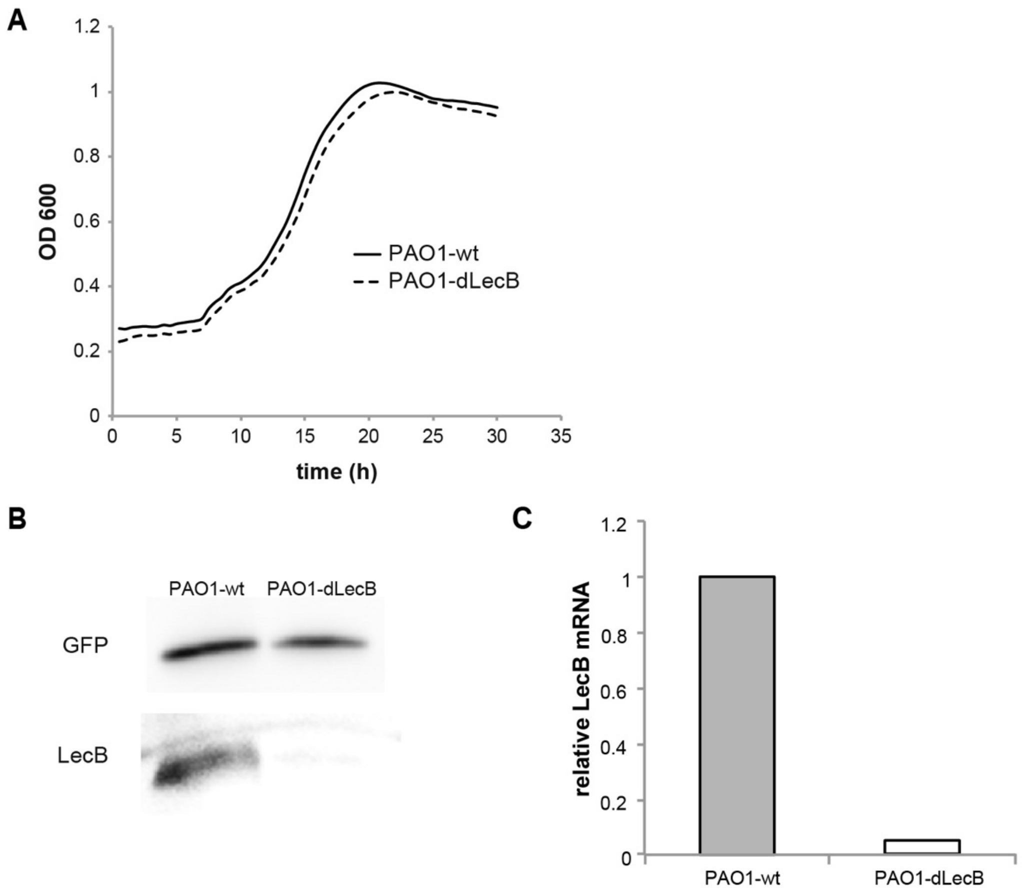

***Figure S7: Control experiments related to Fig. 6***

(A) OD-600 growth curves from wt PAO1 (PAO1-wt) and LecB-knockout strains (PAO1-dLecB) grown in LB medium at 37 °C. (B) – (C) The presence of LecB was probed in PAO1-wt and PAO1-dLecB, both stably expressing GFP, by WB (B) and semi-quantitative qPCR with rpoD as reference (C).

**Table S1: List of basolateral LecB interaction partners identified by SILAC MS**

| # | Protein IDs | Gene name | Number of proteins | Peptides | PEP | log2 (baso vs ctrl) 01 | log2 (baso vs ctrl) 02 |
|---|---|---|---|---|---|---|---|
| 1 | E2QSC8 | TSPAN8 | 1 | 2 | 1,67E-29 | 6,61 | 5,86 |
| 2 | F1PP33;J9NUT8 | ITGA1 | 2 | 12 | 1,38E-73 | 5,97 | 3,14 |
| 3 | F1PR26;J9NW82 | PTGFRN | 2 | 36 | 7,14E-133 | 5,82 | 4,86 |
| 4 | E2RBL9 | ITGB6 | 1 | 12 | 2,41E-46 | 5,74 | 4,82 |
| 5 | E2QZT4 | BSG | 1 | 11 | 2,74E-37 | 5,52 | 5,14 |
| 6 | E2RT60 | ITGB1 | 1 | 18 | 1,23E-132 | 5,47 | 6,33 |
| 7 | J9PB47;A1YV64 | CAECAM1; CEACAM28 | 2 | 4 | 9,72E-38 | 5,36 | 4,52 |
| 8 | O97702;F1PPG5 | GPIIIa;ITGB3 | 2 | 13 | 6,15E-74 | 5,24 | 5,18 |
| 9 | F1PF03;E2RH87 | EGFR | 2 | 16 | 4,79E-76 | 5,22 | 2,85 |
| 10 | E2QX93 | PCDH1 | 1 | 13 | 8,09E-60 | 5,21 | 3,72 |
| 11 | J9P3V6;F1PWJ7 | BCAM | 2 | 20 | 4,82E-239 | 5,21 | 5,26 |
| 12 | F1PRC5 | SLC3A2 | 1 | 16 | 2,43E-64 | 5,09 | 3,91 |
| 13 | B6V8E6 | CTNNB1 | 1 | 15 | 1,43E-82 | 4,98 | 2,99 |
| 14 | F1P9W5;E2RRI1 | SLC12A2 | 2 | 25 | 1,39E-121 | 4,95 | 4,64 |
| 15 | F1P8Q0 | ITGAV | 1 | 33 | 1,48E-213 | 4,88 | 3,72 |
| 16 | F1PUP2;F1PUN5; F1PTF5;F1PTF3; F1PTF2 | PTPRM | 5 | 12 | 1,19E-41 | 4,80 | 2,26 |
| 17 | P33724;P33724-2; F1PWG1 | CAV1 | 3 | 3 | 1,89E-15 | 4,74 | 4,61 |
| 18 | F1PGD5 | LY75 | 1 | 14 | 2,78E-52 | 4,73 | 3,28 |
| 19 | J9NZA9;F1PBA1; P50997;F1P767; F1PL53;F1PJF0; P50996;F1PRH6; J9NZ73;J9P483; F1P8N4 | ATP1A1 | 11 | 36 | 0,00E+00 | 4,71 | 5,06 |
| 20 | E2R9S7;J9NXR7; F1Q015;F1PG43 | CTNNA1 | 4 | 21 | 3,67E-233 | 4,70 | 3,48 |
| 21 | E2RL88;J9NVU0 | ITGA6 | 2 | 26 | 2,35E-149 | 4,69 | 2,99 |
| 22 | E2RMT2;F1PHH3 | PTK7 | 2 | 19 | 1,86E-112 | 4,68 | 3,10 |
| 23 | E2RFE1 | ITGB4 | 1 | 17 | 8,20E-139 | 4,65 | 3,48 |
| 24 | J9NXR3 | Uncharacterized protein | 1 | 13 | 5,23E-63 | 4,58 | 3,46 |
| 25 | F1PTY0;J9PAF7 | SLC4A7 | 2 | 5 | 1,53E-15 | 4,58 | 2,50 |
| 26 | F1PAA3 | CDH3 | 1 | 9 | 3,27E-23 | 4,43 | 4,60 |
| 27 | P31637 | SLC5A3 | 1 | 3 | 1,26E-11 | 4,41 | 2,60 |
| 28 | F1PAA4;F1PAA9 | CDH1 | 2 | 14 | 9,61E-100 | 4,39 | 4,55 |
| 29 | E2R2V6 | L1CAM | 1 | 36 | 1,56E-222 | 4,38 | 3,80 |
| 30 | E2R0Z0;E2R858; J9NXY3;E2R971 | CDH6 | 4 | 16 | 2,27E-49 | 4,37 | 6,21 |
| 31 | F1PV63 | CDCP1 | 1 | 11 | 2,12E-51 | 4,30 | 3,38 |
| 32 | F1P9S5;J9NRH0; J9PA53;E2QY27; E2RDE1 | PLXNB2 | 5 | 27 | 4,97E-140 | 4,24 | 3,04 |
| 33 | F1PXU6;J9NY09; E2RPM8;E2R795 | IGF1R | 4 | 26 | 5,53E-110 | 4,20 | 4,24 |

| # | Protein IDs | Gene name | Number of proteins | Peptides | PEP | log2 (baso vs ctrl) 01 | log2 (baso vs ctrl) 02 |
|---|---|---|---|---|---|---|---|
| 34 | E2REA9 | ITGA2 | 1 | 28 | 1,35E-257 | 4,14 | 4,19 |
| 35 | J9NTL4;F1PK94 | CDH17 | 2 | 16 | 9,47E-99 | 4,07 | 3,42 |
| 36 | J9P423;F1PTZ7;Q28284 | CD44 | 3 | 6 | 1,56E-18 | 3,99 | 6,38 |
| 37 | E2R4F0;J9P8K7 | CELSR2 | 2 | 19 | 1,29E-63 | 3,83 | 2,50 |
| 38 | Q9XSU4;F2Z4N5;J9P7F5 | RPS11 | 3 | 5 | 1,14E-09 | 3,81 | 3,58 |
| 39 | F1PU73 | NCSTN | 1 | 6 | 1,11E-23 | 3,81 | 4,38 |
| 40 | Z4YHE9;Q75ZY9;J9P743;F1PEM3;J9NY14;F1PR52;E2RSH8;F6XRA9 | MET | 8 | 20 | 6,25E-84 | 3,80 | 3,73 |
| 41 | F1Q439 | ITGA3 | 1 | 13 | 3,28E-122 | 3,78 | 5,75 |
| 42 | E2RE80;E2RE81;F1PH17;E2R6Z2 | PTPRF | 4 | 34 | 4,63E-134 | 3,75 | 2,89 |
| 43 | F1PWL4 | PVRL2 | 1 | 5 | 9,06E-24 | 3,71 | 2,54 |
| 44 | F1PGJ7;J9P5N9;E2RNZ4 | DDR1 | 3 | 12 | 2,07E-67 | 3,70 | 4,70 |
| 45 | F1PBI6 | THBS1 | 1 | 11 | 5,93E-43 | 3,67 | 2,54 |
| 46 | F1PRQ8;F1PAD6 | EPHB4 | 2 | 10 | 7,64E-53 | 3,62 | 2,39 |
| 47 | E2QXI6 | TPBG | 1 | 5 | 1,45E-19 | 3,59 | 3,32 |
| 48 | F1PIQ9;O18735;F1PQ05;F1PAC5;J9P9R2;J9P025;E2R7P5;F1P627;E2RHY5;F1P626;J9P9H9 | ERBB2 | 11 | 17 | 9,55E-72 | 3,59 | 4,80 |
| 49 | F1Q1M9;F1PM51 | EPCAM | 2 | 12 | 2,05E-75 | 3,49 | 3,53 |
| 50 | F1PFS1 | STOM | 1 | 5 | 6,91E-32 | 3,28 | 3,62 |
| 51 | J9P897;F2Z4P3;Q9XSU3 | RPL23 | 3 | 4 | 2,21E-12 | 3,27 | 3,29 |
| 52 | P06583;J9P7J0;F1PMF0 | ATP1B1 | 3 | 8 | 3,27E-23 | 3,25 | 4,54 |
| 53 | F1PAF1;E2R7J5 | EPHB2 | 2 | 10 | 1,43E-62 | 3,19 | 3,11 |
| 54 | J9P798;J9P7C2;J9NSL1;E2RPP2;J9JHX3;E2RCJ1;E2R443 | RPS15A | 7 | 5 | 6,47E-18 | 3,12 | 2,22 |
| 55 | F1PPI5 | SLC1A5 | 1 | 5 | 7,18E-40 | 2,88 | 2,82 |
| 56 | E2RH47 | RPS3 | 1 | 8 | 1,60E-55 | 2,85 | 2,53 |
| 57 | E2R9G7;E2QTG3 | RPL11 | 2 | 3 | 1,28E-50 | 2,70 | 2,49 |
| 58 | F1P9K1 | IGSF3 | 1 | 15 | 8,96E-73 | 2,63 | 2,35 |
| 59 | F1PVS7 | PVR | 1 | 8 | 2,09E-27 | 2,50 | 3,56 |
| 60 | F1Q0H9 | SLC1A4 | 1 | 4 | 1,18E-14 | 2,48 | 2,83 |
| 61 | P33729;F1PB95 | ICAM1 | 2 | 7 | 6,33E-27 | 2,44 | 2,25 |
| 62 | F1PRL5;E2RRG4 | SLC25A5 | 2 | 6 | 1,66E-22 | 2,27 | 2,20 |
| 63 | J9P5A2;F1PLY1 | CELSR1 | 2 | 10 | 2,83E-47 | 2,27 | 2,85 |
| 64 | E2R8R8 | RPS9 | 1 | 4 | 4,70E-11 | 2,18 | 2,39 |
| 65 | E2R149 | RPL9 | 1 | 5 | 9,36E-17 | 2,04 | 2,91 |

**Table S2: Lists of used primary and secondary antibodies** (WB … Western Blot, IF … immunofluorescence, IP … immunoprecipitation, surf. stain … used for surface staining in live cells, milk … milk used as blocking agent, methanol … cell fixation with methanol)

*Primary antibodies:*

| antibody target | generated in species | supplier | product number | dilution (application) |
|---|---|---|---|---|
| α3-integrin | mouse | BD Biosciences | 611044 | 1:200 (IF, methanol); 1:1000 (WB) |
| β-actin | mouse | Sigma Aldrich | A5316 | 1:1000 (WB) |
| β-catenin | rabbit | Abcam | ab32572 | 1:250 (IF) |
| β1-integrin | mouse | Thermo Scientific | MA1-19105 | 1:200 (IF); 1:1000 (WB, milk) |
| β1-integrin | goat | R&D Systems | AF1778 | 1:100 (IF) |
| β1-integrin | mouse | Millipore | MAB2000 | 1:200 (IF); 1:1000 (WB, milk); 1:200 (IP) |
| β1-integrin (4B4) | mouse | Beckman Coulter | 6603113 | 1:100 (surf. stain) |
| β1-integrin (9EG7) | rat | BD Biosciences | 550531 | 1:100 (surf. stain) |
| β1-integrin (mAB 13) | rat | BD Biosciences | 552828 | 1:100 (surf. stain) |
| β1-integrin( AIIB2) | rat | Millipore | MABT409 | 1:100 (surf. stain) |
| E-cadherin | mouse | BD Biosciences | 610182 | 1:200 (IF) |
| laminin (antibody detects β1 and γ1 of LM-511 in MDCK cells (*35*)) | rabbit | Sigma Aldrich | L9393 | 1:100 (IF) |
| Lamp1 | rabbit | Cell Signaling | 9091 | 1:100 (IF) |
| LecB | rabbit | Eurogentec | newly produced | 1:1000 (WB) |
| Rab9 | rabbit | Cell Signaling | 5118 | 1:100 (IF) |
| ZO-1 | rat | Millipore | MABT11 | 1:50 (IF) |

*Secondary antibodies:*

| antibody target | conjugation | supplier | product number | dilution (application) |
|---|---|---|---|---|
| anti-mouse | HRP | Cell Signaling | 7076 | 1:2000 (WB) |
| anti-mouse | Alexa488 | Thermo Fisher | A21202 | 1:200 (IF) |
| anti-mouse | Cy3 | Jackson Immunoresearch | 715-166-1500 | 1:200 (IF) |
| anti-mouse | Alexa647 | Thermo Fisher | A21236 | 1:200 (IF) |
| anti-rabbit | HRP | Cell Signaling | 7074 | 1:2000 (WB) |
| anti-rabbit | Alexa488 | Thermo Fisher | A21206 | 1:200 (IF) |
| anti-rabbit | Cy3 | Jackson Immunoresearch | 711-166-152 | 1:200 (IF) |
| anti-rabbit | Alexa647 | Thermo Fisher | A21245 | 1:200 (IF) |
| anti-rat | Alexa488 | Thermo Fisher | A21208 | 1:200 (IF) |
| anti-rat | Alexa647 | Thermo Fisher | A21247 | 1:200 (IF) |
| anti-goat | Alexa647 | Thermo Fisher | A21447 | 1:200 (IF) |
